# Recurrent circuits encode visual center-surround computations in the mouse superior colliculus

**DOI:** 10.1101/2023.09.03.556096

**Authors:** Peng Cui, Kuisong Song, Dimitris Mariatos-Metaxas, Arturo G. Isla, Teresa Femenia, Iakovos Lazaridis, Konstantinos Meletis, Arvind Kumar, Andreas A. Kardamakis

**Affiliations:** Department of Neuroscience, Karolinska Institutet, Stockholm, Sweden; Instituto de Neurociencias CSIC-UMH, San Juan de Alicante, Spain; Tecnológico de Monterrey, Escuela de Medicina y Ciencias de la Salud, México; Department of Neurobiology, Care Sciences and Society, Karolinska Institutet, Sweden; McGovern Institute, Massachusetts Institute of Technology, Cambridge, USA; Division of Computational Science and Technology. School of Electrical Engineering and Computer Science. KTH Royal Institute of Technology, Stockholm, Sweden

**Keywords:** Neural circuit, vision, superior colliculus, surround suppression

## Abstract

Center-surround interactions are fundamental to visual saliency computation, but debate continues over whether and how subcortical visual circuits actively contribute. To address this, we developed an optogenetic approach to delineate the visual center and surround zones of individual neurons in the superficial layer of the superior colliculus (SCs) using only retinal ganglion cell input. Using whole-cell recordings, we demonstrate that surround network activation suppresses center excitability, indicating that SCs circuitry is self-sufficient in driving center-surround dynamics. Through cell-type-specific trans-synaptic tracing and large-scale modeling, we identified an SCs-based circuit with two key motifs driving surround modulation: recurrent excitation and feedback inhibition. We propose that subcortical visual circuits in the SCs have evolved to perform surround suppression alongside retinal and cortical suppression, facilitating the distribution of parallel saliency computations across different levels.

**Significance statement:** This study questions the notion that the superior colliculus (SC) merely acts as a passive recipient of saliency information from upstream circuits. We demonstrate that the SC can independently generate center-surround interactions that could contribute to visual saliency through local circuits without top-down input. This ability represents a computation that has been conserved since the dawn of vertebrate evolution. By mapping these interactions, we reveal that the mouse SC actively induces visual surround suppression. These findings suggest that phylogenetically older circuits in the SC may play a more independent role in active vision than previously acknowledged, prompting a reevaluation of visual saliency processing across subcortical brain regions.

## Introduction

Enabling neurons to compare inputs to their receptive field with their immediate surround represents a fundamental and conserved computation (1, 2). This mechanism is widespread across sensory modalities, including vision (3), hearing (4), somatosensation (5), and olfaction (6), and is typically suppressive, a process known as ‘surround modulation’. A typical example found across species is during size tuning of a visual neuron; the firing response initially increases when a stimulus is centered on its receptive field (7, 8), but then decreases as the stimulus expands beyond this area (9–12). Such observations have been measured throughout the visual thalamocortical system in the lateral geniculate nucleus (13), visual cortex (14) and extrastriate cortex (15), as well as in retinotectal systems in the premammalian optic tectum (9, 16, 17) and superior colliculus (8, 10).

While center-surround interactions are known to originate in the retina (18–21), the extent of their modulatory effects in later stages of the visual hierarchy, such as in the superior colliculus (SC) and visual cortex (VIS), and their interrelationships remain unclear (Fig. 1). In this study, we investigated whether local circuits within the superficial layers of the superior colliculus (SCs) can perform center-surround interactions using solely retinal input, independent of visual cortical or other extrinsic inputs. Understanding this computation is crucial for elucidating how top-down cortical signals in turn interact with SC networks in saliency processing (22–24) and in visuospatial behaviors, including target selection during overt (25, 26) and covert attention (27, 28)

**Fig. 1.**
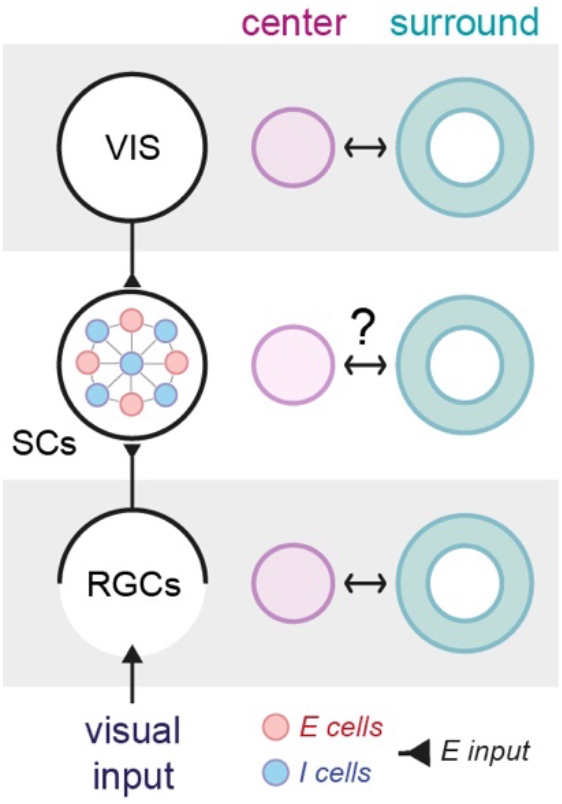
*(Left)* Schematic illustrating the two primary sources of glutamatergic visual input (*E input*) targeting excitatory (*E cells*, red) and inhibitory neurons (*I cells*, blue) in the superficial layers of the superior colliculus (SCs), originating directly from retinal ganglion cells (RGCs) and indirectly from the visual cortex (VIS). *(Right)* Visual center and surround interactions are well understood in the retina and visual cortex. However, it remains unclear whether the observed surround modulation in the SCs (8, 10) is simply conveyed from suppression already occurring in the retina and visual cortex, or if the SCs can actively generate center-surround interactions.

We developed a method that combines optogenetics and whole-cell electrophysiology to map the center and surround zones of individual SC neurons in midbrain slices. Here, we delineate their center excitation zones in response to inputs provided by RGCs. By selectively activating these zones in sequence, we reveal that the mouse SCs contains the circuitry essential for surround suppression, a mechanism conserved in vertebrate evolution (9). Moreover, like in the visual cortex (29), we found that surround suppression in SCs neurons is primarily driven by decreases in excitation rather than increases in synaptic inhibition, as classically expected. To establish its circuit basis, we performed cell-type-specific trans-synaptic mapping to identify and quantify the connectivity patterns defining the circuit motifs between local excitatory (*E*) and inhibitory (*I*) neurons in the SCs. Large-scale modeling of these connections revealed two key motifs driving surround suppression in the SCs: recurrent excitatory (*E-E*) and feedback inhibitory (*I-E*). This finding aligns with model predictions involving strong recurrent networks in the visual cortex and their role in cortical visual suppression (30, 31), despite manifesting in a structure that predates the evolution of the neocortex.

## Results

### Mapping retinal excitation zones of single SCs neurons

By leveraging the aligned topographic correspondence of the retinotectal pathway (32), we generated an input-dependent visual field and made it light-excitable by injecting an adeno-associated virus (AAV) intravitreally that expressed ChR2 in retinal ganglion cells (RGCs) of both wild-type (*n =* 22, *N =* 14) and vGAT-Cre (*n =* 7, *N =* 5) mice (Fig. S1A). For trial inclusion, we first confirmed sufficient expression of ChR2 across the SCs and homogeneous pan-retinal expression of ChR2 (Fig. 2A, Left); partial expressions were not taken into consideration to minimize the risk of false positive surround classification that could arise from lack of ChR2 expression. Retinal excitation zones of single SCs neurons (*n =* 29, *N =* 19) were subsequently mapped out by delivering retinotopic light-impulse patterns to discrete sub-regions of the coronal SCs (100 × 70 μm each) while recording excitatory postsynaptic currents (EPSCs) in whole-cell configuration (Fig. 2B and Fig. S1).

**Fig. 2.**
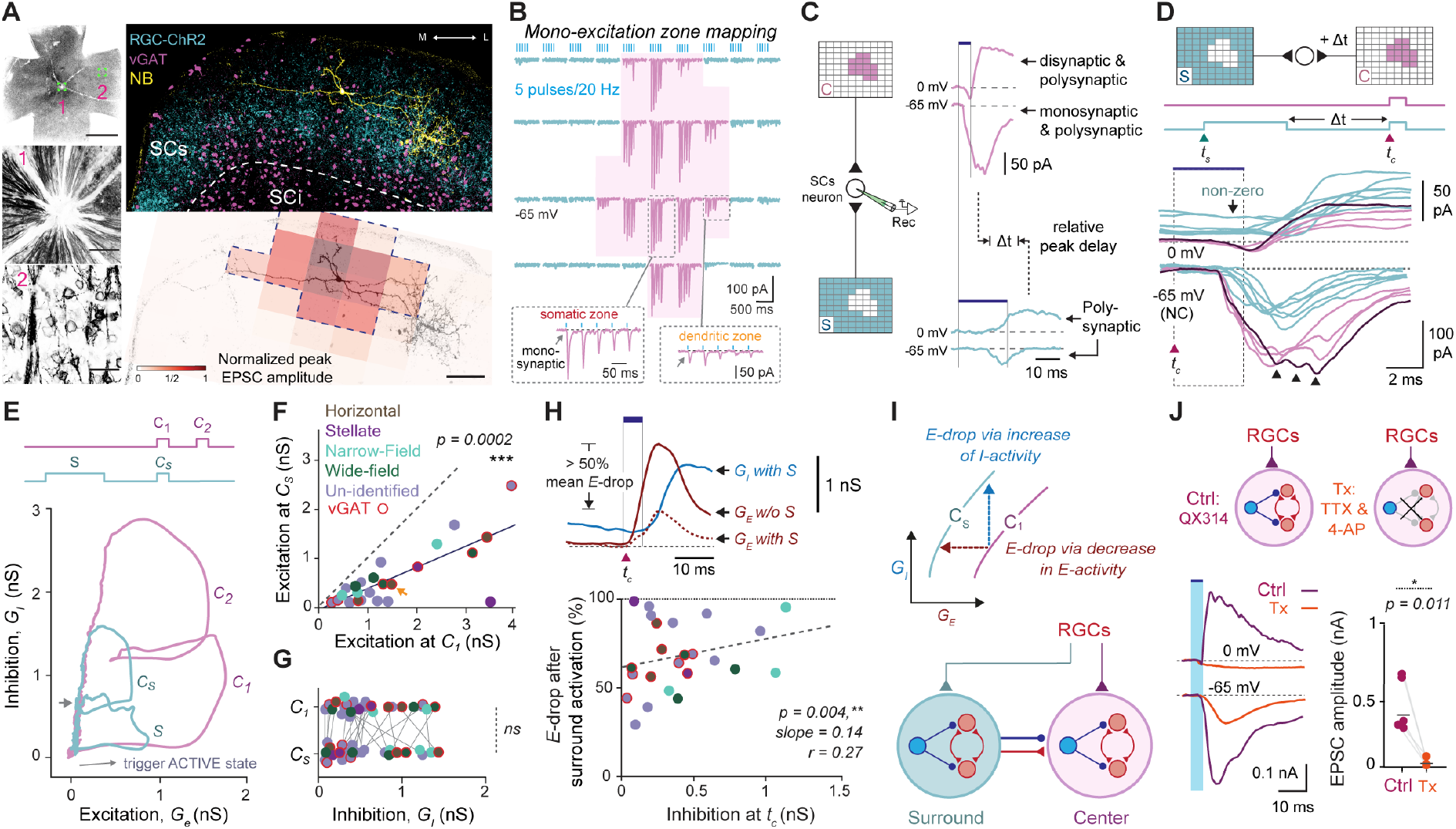
Visual surround network activation modulates center excitability in the SCs. *(A, Left)* From top to bottom: Inverted fluorescent image of whole retinal mount showing homogeneous transfection of RGCs with ChR2-EYFP. Scale bar: 1200 μm. Images below show enlarged aspects of the optic disc (1) and individual RGCs (2). Scale bar: 25 μm. Bottom right: Coronal midbrain section with ChR2-expressing RGC axonal terminals seen occupying the SCs layers. (*Right*) Top: A coronal section of the SCs that shows a representative example of this approach applied to an inhibitory horizontal SCs neuron (vGAT+, magenta) stained intracellularly with Neurobiotin (NB, yellow). ChR2-expressing RGC axonal terminals are shown in cyan. Bottom: Reconstructed neuron is shown with the denoted excitation zone (dotted line), which is referred to as center, whereas the non-responsive areas as surround (S) zone by mapping its monosynaptic excitation zone (shown in B). *(B)* Excitation zone mapping applied in neuron in (A) used for center zone determination. Light pulse trains (20 Hz, 5 ms) trigger excitatory postsynaptic currents (EPSCs, shown in magenta) with fixed short-latency responses suggesting monosynaptic transmission. Each trace is evoked from a single optostimulated rectangular area of 100 × 70 μm. When the rectangle is positioned over somatic zones, the responses are larger compared to those from dendritic zones; together, they delineate the overall center response zone of the neuron. *(C, Left)* Schematic showing two different light patterns used for optostimulation of ChR2-expressing RGC axonal terminals in the center (top) and surround (bottom) zones, while whole-cell recording postsynaptic responses from an SCs neuron. *(Top right)* Optostimulation of the center zone evoked excitatory and inhibitory postsynaptic currents when recording near reversal equilibriums for GABA_A_-chloride mediated inhibition and glutamatergic excitation (−65 and 0 mV, respectively). *(Bottom right)* Surround network activation triggers long-latency polysynaptic excitatory and inhibitory activity. The relative delay between peak center excitatory and peak surround inhibitory, *Δt*, was on average 20 ms. Traces were not corrected (NC) for the offset created by the liquid junction potential, which was experimentally confirmed to be ∼10 mV. *(D)* Synaptic modulation between the center and surround zone of the neuron shown in (a–c). Top: Surround and center interaction performed in a collision-style fashion by introducing delay *Δt* between surround and center stimulation. (Bottom) Individual traces of synaptic inward and outward currents evoked in response to center stimulation Arrowheads show oligosynaptic or recurrent activity (darker shade). Scale bar: 100 μm. Full traces in Fig. S2. *(E, Top)* Optostimulation patterns for paired-pulse center-only and single-pulse transition surround-to-center activation. *(Bottom)* Plot of the estimated inhibitory conductance as a function of the excitatory conductance in the neuron shown in (a-d). Arrow shows direction of process when neuron is activated. Smaller gray arrow indicates point of center stimulation in the surround-to-center case. *(F*–*G)* Comparison of estimated synaptic excitatory (F) and inhibitory (G) conductances between two conditions: center-only (C_1_) and center after surround (C_s_). Color coding corresponds to morphologically or genetically identified SCs neuron cell-types. Orange arrow depicts data point from the neuron shown in (a– d). *(H, Top)* Estimated excitatory conductance triggered by center-only (C_1_; solid red line) and center-surround (C_s_; dashed red line) condition. Inhibitory conductance (blue) in the center-surround condition; note surround inhibition is non-zero at time *t*_*c*_. *See* Fig. S3 for all traces. *(Bottom)* Modulation of center excitation plotted as a function of surround inhibition at *t*_*c*_. *(I, Top)* Schematic showing two theoretical lines indicating iso-conductance curves relating excitation to inhibition. Response suppression involves transitioning from a higher level to a lower level curve. The cartoon shows how the same level of response suppression can be achieved by either decrease in excitatory conductance or by increase in the inhibitory conductance. *(Bottom)* Circuit diagram illustrating the direct and indirect actions of surround inhibition in the SCs through connections between excitatory (red) and inhibitory (blue) neurons in the center and surround. Note the recurrent connections between excitatory neurons. *(J)* Excitatory and inhibitory current recordings (at -65 and 0 mV) from SCs neurons (*n =* 5) were performed in response to center optostimulation of ChR2-expressing RGC axonal terminals. Bath application of 1 μM TTX and 100 μM 4-AP isolate monosynaptic RGC-triggered excitation while simultaneously abolishing recurrent excitation and recruited inhibition. Statistical comparisons were performed as paired *t*-tests followed by Wilcoxon rank-signed tests. *Abbreviations: NC*, not corrected.

Figure 2A-B illustrates the logic of this approach by showing the center zone of a vGAT+ horizontal cell, identified by detecting regions with fixed short-latency EPSCs (held at -65 mV) indicative of monosynaptic input following optostimulation (5 pulses at 20 Hz) of a 10 × 10 grid spanning the SCs (Fig. 2B). The areas that evoked EPSC amplitudes exceeding >∼10% of the peak value (Fig. S1E–F) were selected, which collectively formed an image pattern that was classified as the center and usually encompassed the soma and most of the dendritic processes (Fig. 2A, Bottom right; *see* also bottom insets in Fig. 2B). Using a large field of view (16×; *see* Fig. S1B), we systematically accounted for the variable spatial extent of a given neuron’s center zone and effectively defined its boundaries in relation to the surrounding region within the SCs (Fig. S4D).

### Retinal-driven surround network activation modulates center excitability in SCs neurons

Selective activation of the entire center zone consistently triggered short-latency compound EPSCs (120.8 ± 15.5 pA at -65 mV; *see* magenta traces in Fig. 2C), often with prominent multimodal peaks (Fig. 2D, *see* arrowheads), suggesting oligosynaptic input and/or recurrent activity in addition to monosynaptic excitatory RGC input. In line with previous studies (9, 33), we observed a di-/oligosynaptic inhibitory postsynaptic current (IPSC; mean amplitude ± s.e.m.: 24.3 ± 5.0 pA at 0 mV) near the equilibrium potential for glutamate-mediated excitation (magenta traces in Fig. 2C–D). Since surround network activation is not monosynaptic and requires interneuronal recruitment, evoked responses were typically long-latency and polysynaptic (cyan traces in Fig. 2C). Latency differences in signaling between center and surround was prominent, where surround excitation and inhibition peaked at average delays of 20 and 28 ms, whereas center excitation and inhibition peaked at 9 and 15 ms, respectively (Fig. S3D).

We induced center-surround interactions in a collision-style manner by introducing a relative delay (Δ*t*) to account for the surround response latencies (Fig. 2D). The surround stimulation phases lasted for 20 ms and were terminated 20 ms before the onset of center stimulation (*t*_*c*_) to ensure adequate time for the buildup of surround-induced activity while synchronizing with the center-induced activity (*see* Fig. S2 and Methods for tested range of values). At *t*_*c*_, only the surround-induced inhibitory components persisted, while any remaining excitation was quenched due to the relatively faster transmission of synaptic excitation (cyan traces in Fig. 2D). This is indicated by the non-null values of surround-triggered outward currents at 0 mV (*see* arrow in Fig. 2D). In this case, the comparison of the evoked EPSCs (at -65 mV; magenta and cyan traces in Fig. 2D) illustrates a consistent decline (> 50% in amplitude in this cell) in center excitability following surround activity.

To provide a more comprehensive view on the mode of surround suppression, we calculated the theoretical contributions of excitatory (*G*_*e*_) and inhibitory (*G*_*i*_) synaptic conductances (34, 35)(*see* also Methods). Figure 2E shows the relationship between inhibition and excitation of the recorded neuron plotted in Fig. 2D during center only (magenta) and center-surround stimulation (cyan; *see* also Fig. S3A). The neuron was subjected to paired-pulse center-only stimulation (C_1_-C_2_), demonstrating that inhibition levels nearly doubled compared to the first pulse (C_1_), while peak excitation was maintained stable after each pulse. During center-surround stimulation, however, inhibition did not significantly increase, despite the neuron’s capacity for higher inhibition levels. Notably, peak excitation was halved (compare C_s_ to C_1_; *see* gray arrow in Fig. 2E), indicating a reduction in excitation without an expected rise in inhibition, suggesting that surround networks act both directly and indirectly to reduce center excitability.

The degree of surround modulation of center excitability varied across different recorded SC neurons, but it consistently resulted in suppression regardless of cell type. More than half of the neurons could be classified based on morphology (e.g., horizontal, wide-field, narrow-field, or stellate; for details, *see* (36)) or genotype (e.g., vGAT+), as quantified in Fig. S4. We categorized the suppression as mild, moderate, or strong, based on the magnitude of the decrease in center excitatory conductance (Fig. 2F): strong (> 60%, *n* = 9/24), moderate (20– 60%, *n* = 10/24), and mild (< 20%, *n* = 5/24). Overall, peak center excitation (Fig. 2F; *G*_*e*_: mean amplitudes of C_1_: 1.7 ± 0.3 nS and C_s_: 0.6 ± 0.1 nS, with a mean percentage decrease of 62.3 ± 4.7%, *p = 0*.*0002*) and peak total conductance (i.e., *G*_*t*_ = *G*_*e*_ + *G*_*i*_: mean amplitudes of C_1_: 2.1 ± 0.3 nS and C_s_: 1.2 ± 0.2 nS, with an average percentage decrease of 40.6 ± 5.3%, *p = 0*.*0007*; *see* Fig. S3B) of the center response were significantly reduced (p = 0.002 an) following surround network activation. However, no significant increase in center inhibition *G*_*i*_ was observed at peak *G*_*t*_ (Fig. 2G; mean amplitudes of C_1_: 0.53 ± 0.08 nS and C_s_: 0.55 ± 0.09 nS). Pharmacological blockade of GABA_A_ receptors with bath-applied gabazine (GBZ) eliminated the inhibitory effect of surround input, transforming responses into seizure-like activity and saturating center response profiles (bottom traces in Fig. S2A). Overall, surround suppression was observed across all SC cell types, regardless of morphology or genotype, with surround inhibition driving decreases in excitation levels.

### Direct and indirect effects of surround inhibition in the SCs

There are two potential explanations for the observed weakening of center excitatory conductance: (i) a direct effect of surround inhibition, which may operate either linearly or non-linearly, and (ii) an indirect effect of surround inhibition, which may reduce the number of active inputs from local excitatory interneurons. To assess the first explanation, we examined the relationship between the inhibitory conductance at *t*_*c*_ (mean amplitude ± s.e.m.: 0.46 ± 0.08 nS, *p* = 0.004, *n* = 24) and the relative drop in peak center excitation (Fig. 2H). This revealed a statistically significant correlation yet weak slope value, indicating that direct effects can only partially explain the observed decrease in center excitation. Therefore, the second explanation need be true to fully explain the excitation drop with indirect surround inhibition acting on other center excitatory neurons, disrupting recurrent connections that would normally enhance excitation thereby curbing responsiveness to center stimuli.

Figure 2I (Top) illustrates two hypothetical iso-conductance curves that relate input excitation and inhibition, where response suppression corresponds to a transition from one curve to the other. The schematic demonstrates that this can be achieved by either decreasing excitatory conductance or increasing inhibitory conductance, or any vector in between. Figure 2I (Bottom) illustrates how this transition could be implemented, as center and surround zones reciprocally influence each other through synergistic putative recurrent interactions among local populations of excitatory and inhibitory neurons.

To explore the efficacy of such a recurrent excitatory connection (*E-E)*, we patch recorded SC neurons (*n = 5*) in response to optostimulation of RGC terminals within their identified center zones in the presence of 1 μM TTX and 100 μM 4-AP (washout *not shown* for clarity). Evoked EPSC amplitudes were drastically reduced with a small component persisting resulting from monosynaptic RGC input (Fig. 2J, mean amplitudes ± s.e.m.: for control: 405.5 ± 81.1 pA; for Tx: 22.3 ± 11.0 pA). This observation demonstrates that a functional *E-E* connection could provide additional center-dependent excitatory input, amplifying incoming retinal signals targeting the center zone, while the absence of outward currents during treatment (held at 0 mV) suggests the removal of recruited di- and/or oligo-synaptic inhibition, revealing the indirect functional impact of feedback or recruited inhibition.

### SCs space defined by recurrent excitatory and inhibitory circuit motifs

To determine whether collicular surround inhibition relies on direct connections between surround inhibitory neurons and center neurons, indirect connections through inhibitory center neurons activated by excitatory surround neurons, or a combination of both, we performed retrograde cell-type-specific trans-synaptic mapping of excitatory (vGluT2+) and inhibitory (vGAT+) neurons. Despite the similar proportions and intermingling of cell types within the SC (37)—which complicates the identification of their connectivity patterns and functions (Fig. 3A)—we were able to differentiate local monosynaptic and presynaptic sources of input based on their neurotransmitter type.

**Fig. 3.**
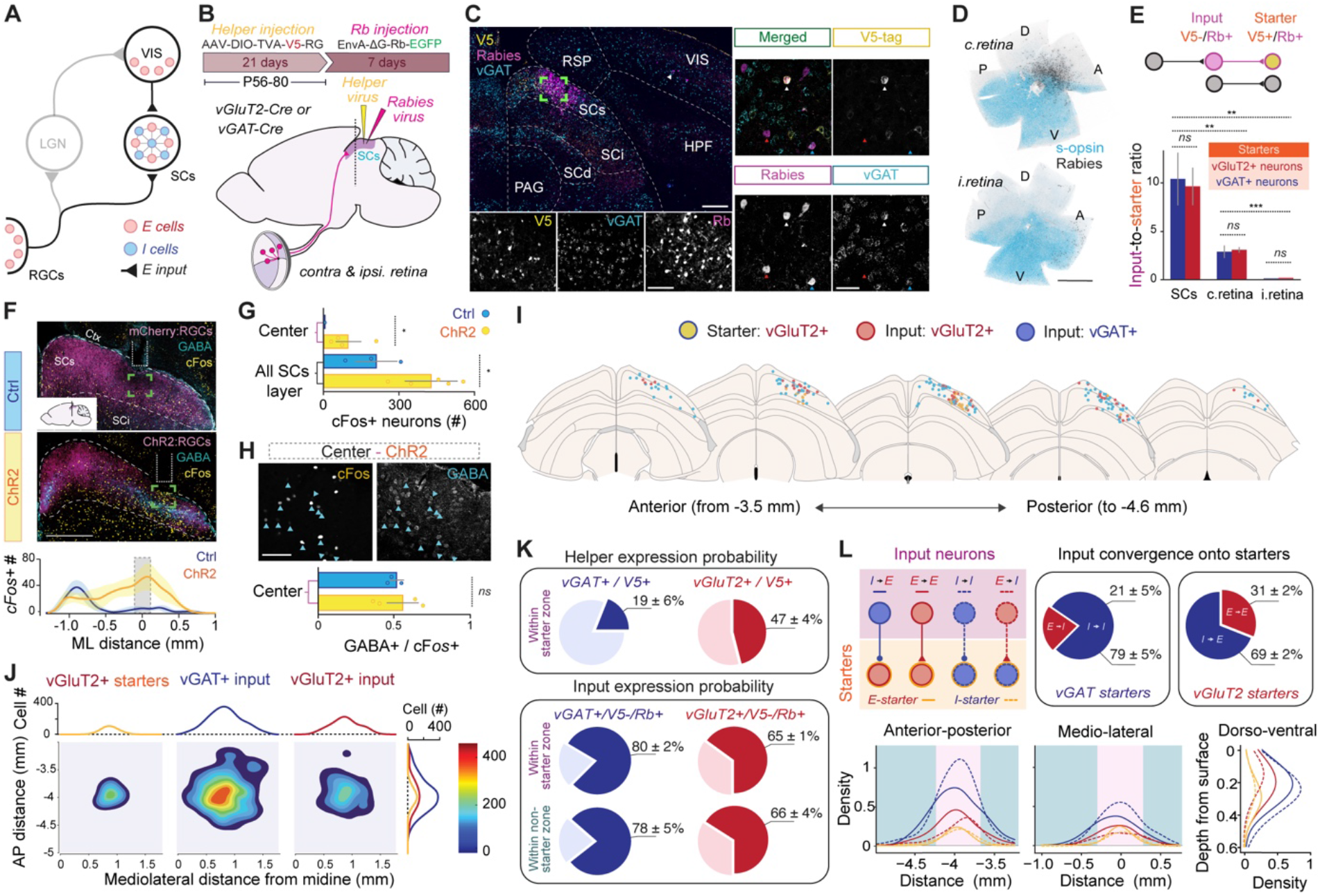
Connectivity logic between local excitatory and inhibitory neurons in SCs space. (*A*) Connectivity diagram of the early visual system highlighting the main sources of visual excitation arising from retinal ganglion cells (RGCs) and visual cortical (VIS) input targeting excitatory and inhibitory neurons in the SCs. (*B*) The helper virus AAV-DIO-TVA-V5-RG expresses the rabies glycoprotein (RG), and a fusion of the TVA receptor with the V5 tag. The RG-deleted rabies virus Rb, pseudotyped with EnvA, expresses EGFP. The AAV helper virus was first injected into vGluT2-Cre (*N =* 3) and vGAT-Cre (*N =* 3) mice and 3 weeks later the Rb-EGFP was delivered in the same location in the SCs. (*C*) Left: coronal section of the midbrain of a vGAT-Cre animal with emphasis on initial injection site where expression of the V5 tag (in yellow) should be restricted to and not beyond the SCs layer. Subsequently, EGFP-expressing neurons are a result of Rb-transfection (magenta). Scale bar: 300 μm. Insets below show colocalization of the enlarged area (green box). Scale bar: 100 μm. Right: RNAscope *in situ* hybridization for *vGAT* (for *vGluT2, see* Fig. S5) revealed the identity of Rb-EGFP input neurons while confirming starter neuron cell-types. White arrows indicate starter neurons, whereas red and blue show excitatory and inhibitory input neurons, respectively. Scale bars: 50 μm. (*D*) Contralateral and ipsilateral retinas showing retinotopy of presynaptic Rb-EGFP RGCs (black). Counterstaining with s-opsin (light blue) reveals retinal orientation. Dorsal, D; anterior, A; ventral, V; posterior, P. Scale bars: 1000 μm. *(E, Top)* Diagram showing how starter neurons are identified based on the co-expression of the V5 tag and Rb-EGFP. Input neurons only express Rb-EGFP and provide monosynaptic and presynaptic input to the starter neurons. *(Bottom)* Input convergence shows the ratio of input-to-starter neurons, i.e., N(input)/N(starter), arising from: i) within the SCs, ii) contralateral (c.), and iii) ipsilateral (i.) retina. *(F, Top)* Coronal midbrain sections of the SCs centered around the implant for *cFos* induction following local optostimulation. Yellow, *cFos* immunostaining; magenta, control (top) or ChR2 (bottom); cyan, GABA immunostaining. Scale bar: 500 μm. *(Bottom)* The location (top) and distribution (bottom) of *cFos*+ neurons across the hemispheres of the SCs. Blue circles are control; Yellow circles are ChR2-induced. The central zone (C) is defined by the implant tip, which defines the origin of the x-axis. Shading is standard error. (*G*) Comparison of the total cell count of *cFos*+ cells induced in SCs neurons between control (Ctrl, blue; *N = 3*) and during activation of ChR2-expressing RGC axonal terminals (ChR2, yellow; *N = 5*) across the entire SCs layer and center zone, respectively. Comparison of means performed using a *t*-test. *(H, Top)* Grayscale image taken from the center region showing *cFos* (left) and GABA immunostaining (right). Blue arrows show colocalization. *(Bottom)* Ratio of GABA+ to *cFos*+ cells compared between the two conditions (ChR2, *N* = 4; and Ctrl, *N* = 3). Statistical testing was done using *t*-test (*, *p* < 0.05; ***, *p* < 0.0001; ns, no significance). Bars show standard error of the mean. (*I*) Schematic depicting the dorsoventral and mediolateral location of starter and input neurons in five separate anterior-posterior (AP) coronal sections of the SCs layer in a vGluT2-Cre animal. *(J)* Contour plot showing the spatial distribution of *E* and *I* input neurons to *I* starter neurons in the AP and medio-lateral (ML) dimensions across the SCs (for case shown in I). *(K, Top)* Helper expression probability showing the percentage of neurons expressing V5 out all the vGAT (left) and vGluT2 (right) neurons located within starter zone. *(Bottom)* Input expression probability for both vGAT and vGluT2 neurons inside and outside the starter zone. The probability is calculated on an *N = 3* for each genotype. *(L, Top left)* Schematic of all possible motifs between input and starter neurons. *(Top right)* Quantitative cell-type-specific input-to-starter ratios (referred to as *input convergence*) reveal feedforward (i.e., *E-I* and *I-E*) and recurrent connectivity patterns (i.e., *E-E* and *I-I*) calculated by *n*(vGAT+)/*n*(Total) and *n*(vGluT2+) / *n*(Total) presynaptic input neurons. Comparison of means performed using a *t*-test. *(Bottom)* Quantitative relationship between the density of input neurons and starter neurons for *E* and *I* cell types across the three anatomical dimensions (AP, ML and DV). Solid lines depict data obtained from excitatory starters averaged from vGluT2-Cre animals (*N* = 3), whereas dashed lines show data obtained from inhibitory starters averaged from vGAT-Cre animals (*N* = 3). *See* Fig. S5E for the distributions obtained from individual animals. Magenta shaded area depicts starter zone; cyan-shaded area is outside the starter zone. *Abbreviations: c*., contralateral; *i*., ipsilateral; *RGCs*, retinal ganglion cells; *LGN*, lateral geniculate nucleus; *VIS* (and *V1*), primary visual cortex; *SCs*, superior colliculus superficial layer; *SCi*, superior colliculus intermediate layer; *SCd*, superior colliculus deep layer; *RSP*, retrosplenial cortex; *PAG*, periaqueductal gray matter; *HPF*, hippocampal formation.

To trace local afferent inputs to vGluT2+ and vGAT+ SC neurons, we microinjected an AAV-DIO-TVA-V5-RG into the SCs of adult vGluT2- and vGAT-Cre mice, using a Cre-dependent AAV helper vector to express both the TVA receptor and rabies glycoprotein (RG) specifically in excitatory or inhibitory neurons (Fig. 3B). Three weeks later, an EnvA-coated, G-deleted rabies (Rb) viral vector expressing enhanced green fluorescent protein (EGFP) was injected at the same site (Fig. 3B–C). Neurons co-expressing Rb and the V5 tag, identified via V5 immunohistochemistry, were labeled as ‘starter’ neurons (Top, Fig. 1E). In vGluT2-Cre animals (*N* = 3), excitatory cells served as starters, while in vGAT-Cre animals (*N* = 3), inhibitory cells were the starters, whereas neurons labeled only with Rb-EGFP were classified as ‘input’ neurons, which provide monosynaptic input to the starter neurons (Fig. 3C–E).

Instead of mapping long-range inputs to the SCs (*see* (38)), we focused on the local connectivity between vGluT2+ and vGAT+ neurons within the SCs, along with the long-range input from RGCs. The low injection volume (50 nL, 1×10^12^ vg/mL) allowed for localized expression of the helper virus within SCs boundaries, without spillover to the intermediate layer (SCi). This also maintained a low infection rate (19% for vGAT+ and 43% for vGluT2+; *see* Fig. 3K, top) allowing for sparse labeling of starter neurons, thereby increasing confidence in quantifying true input neurons (V5-/Rb+) within the starter zone, defined by co-labeled V5+/Rb+ starter neurons. The similar input expression probabilities inside and outside the starter zone (80% and 78% for vGAT+ and 65% and 66% for vGluT2+, respectively; *see* Fig. 3K, bottom) further validated accurate assessment of input neurons within the starter zone.

We first quantified the total number of presynaptic SCs input neurons (Fig. 3E) along with long-range inputs from both contralateral and ipsilateral retina (Fig. 3D). To determine whether they were excitatory or inhibitory, we performed RNAscope *in situ* hybridization with *vGluT2* or *vGAT probe*, respectively (*see* Fig. 3C and Fig. S5). Co-staining of Rb-EGFP+ cells with *vGluT2* and *vGAT* probes revealed no colocalization, as neurons negative for *vGAT* were positive for *vGluT2* and vice versa, with nearly all detected input cells labeled by only one of the two probes (269 out of 272 in *N* = 2 control animals; Fig. S5). Inputs to vGluT2+ and vGAT+ starter neurons from SC interneurons were comparable, averaging 9.5 ± 2 and 10.2 ± 2.8 cells, respectively. The retinotopic pattern was evident, with contralateral RGC inputs primarily originating from neurons in the central regions of the retina and ipsilateral RGC inputs localized to peripheral regions (Fig. 3D), while both types of RGCs contacted a balanced number of excitatory and inhibitory starter SC neurons (Fig. 3E; 2.9 ± 0.3 for vGluT2+ and 2.8 ± 0.7 for vGAT+ for contralateral RGCs; 0.7 ± 0.01 for vGluT2+ and 0.3 ± 0.02 for vGAT+ for ipsilateral RGCs).

### Retinal input recruits balanced excitatory and inhibitory neurons

To assess whether retinal input functionally recruits excitatory and inhibitory inputs - as suggested in Fig. 3E - or if there is selective recruitment, we performed intravitreal injections in wild-type mice using either AAV2-ChR2-EYFP (*N* = 5) for ChR2 expression or AAV2-mCherry (*N* = 3) as a control, targeting RGC axonal terminals (Fig. 3F and Fig. S6). Unilateral optostimulation of RGC terminals was performed using an implanted fiber optic onto the dorsal aspect of the SCs, and neuronal activation was confirmed through *cFos* labeling, a surrogate marker of neural activity (Fig. 3F and Fig. S6; *see* Methods). Figure 3F (Bottom) shows the distribution of active SC cells, highlighting a significant increase in *cFos* expression under the implant in RGC-ChR2 mice compared to the control (Fig. 3G). Figure 3H shows a subset of *cFos* positive neurons in the center zone that were immunoreactive for the neurotransmitter GABA revealing their inhibitory origin (*see* also Fig. S7). Under these experimental settings, RGC input drives a balanced recruitment of excitation and inhibition in the center with a population of putative glutamatergic neurons (i.e., negative for GABA) that closely matches the population size of active inhibitory neurons (ratio: 0.49 ± 0.07, *N =* 4; Fig. 3H). This result suggests that a visual stimulus, whether center or surround, activates both cell populations in similar proportions.

### Every connectivity pattern is possible in the SCs

Figure 3I illustrates the spatial distributions of vGluT2+ starter neurons and vGluT2+ and vGAT+ input neurons, mapped by detecting the cells expressing V5 and/or RV-EGFP and locating them within midbrain sections from the Allen Reference Atlas coordinate system. Gaussian kernel-based density plots in Fig. 3J reveal their horizontal distribution patterns along the ML and AP dimensions (Fig. S5E). In this vGlut2 animal, excitatory starter neurons (left panel in Fig. 3J) occupied a confined retinotopic area within the SCs, influenced monosynaptically by a broader excitatory area (right panel), as well as an even broader inhibitory area (middle panel).

Using this approach applied in all animals (*N =* 6; 3 vGluT2 and 3 vGAT mice), we found that all possible connectivity patterns between input and starter neurons existed (Fig. 3L). Color-coded solid lines represent average spatial distributions using excitatory neurons as starters, while color-coded dashed lines correspond to those from inhibitory starters (for each animal, *see* Fig. S5E). Input neurons always displayed a higher normalized density distribution compared to their starter counterparts. A stable E/I ratio is maintained throughout the SCs with *I-E* and *I-I* input-to-starter connections spanning broader ranges on all ML, AP and DV axes. In contrast to excitatory input, inhibitory inputs formed a significantly larger number of connections with inhibitory starter neurons (78.8 ± 4.6%, *n* = 3) compared to the 21.2 ± 4.6% (*n = 3*) established by excitatory inputs. Similarly, inhibitory inputs connected with excitatory starter neurons at a higher rate (69.2 ± 2.3%, *n* = 3) compared to the 30.8 ± 2.3% (*n = 3*) connections made by excitatory inputs. Approximately one-third of excitatory input neurons establish monosynaptic contacts with excitatory starter neurons, thereby forming a recurrent excitatory network that can be accessed monosynaptically by functional RGC input (Fig. 2J).

### Recurrent excitation (*E-E*) and feedback inhibition (*I-E*) necessary for surround suppression

We sought to identify the minimal connection or combination of connections needed to account for the observed electrophysiological (Fig. 2) and circuit mapping (Fig. 3) results. The summarizing cartoon in Fig. 4A depicts how the spatial distribution and input convergence ratios (Fig. 3L) can delineate the SCs circuit into a hypothetical visual center and a surround. To evaluate the contribution of each identified connection in Fig. 1 (i.e., *E-E, E-I, I-I*, and *I-E*), we developed a large-scale model based on Fig. 4A, consisting of 12,800 spiking neurons (*see* Table S3 for stimulus input; Table S4 for neuronal parameters; Table S5 for synaptic parameters; Table S6 for network connectivity; and Table S7 for external input parameters). This model allowed us to explore connection strengths and simulate interactions between excitatory and inhibitory SC neurons and isolate the critical ones contributing to SCs surround suppression.

**Fig. 4.**
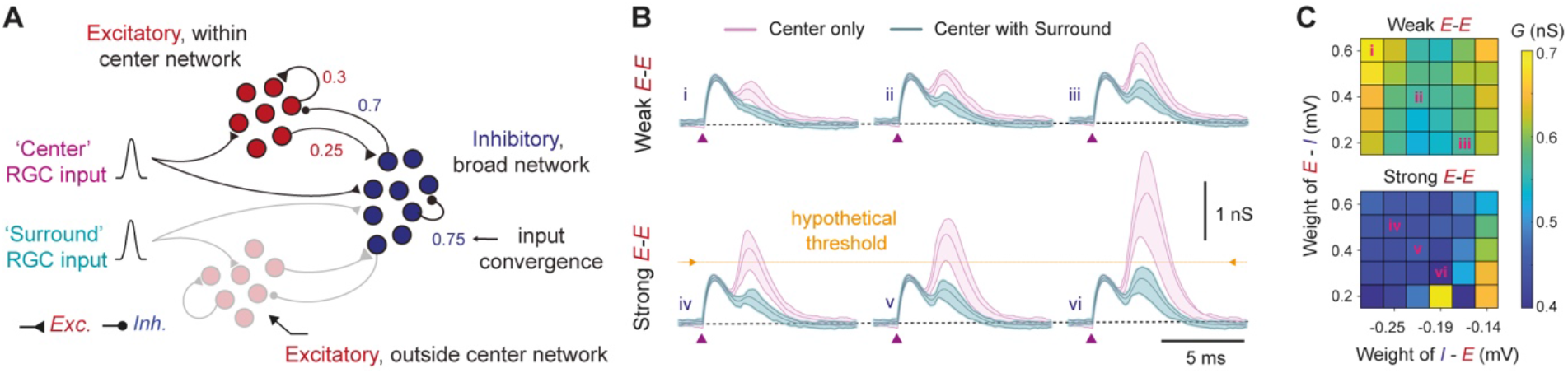
Model predicts recurrent networks a key feature in visual surround modulation. (*A*) Schematic of the E/I connectivity in SCs network highlighting the concept between center and surround in context of retinal input. Network model was simulated based on spatial distribution and input convergence using 6400 excitatory and 6400 inhibitory neurons. Raster of spiking activity and spatial arrangement of center and surround network response profiles are shown in Fig. S7. (*B*) Time courses of excitatory conductance of a center neuron. Each trace is an average of 20 trials and shading indicated 25%–75% quartiles. The first peak is generated with low-variance monosynaptic activation of direct retinal input whereas the second peak is a network effect generated by balanced recurrent excitation and recurrent inhibition. The connectivity parameters for these six examples are marked in inset (C) with Latin numerals. (*C*) Normalized center conductance during surround stimulation. Each colormap shows normalized change in excitatory conductance (*see* Methods) in a center neuron (shown in *B*) as a function of recurrent excitation (top: weak connection; bottom: strong connection) and mutual connectivity weights between excitation and inhibition (x-axis: inhibition of excitatory neurons; y-axis: excitation of inhibitory neurons).

To mimic center zone activation, we triggered activity in 314 excitatory and 314 inhibitory model neurons and measured the evoked excitatory synaptic conductance in a randomly chosen neuron from the center region (Fig. 4B, *see* Fig. S7). First, the center neurons alone were stimulated (magenta traces in Fig. 4B). The first peak was provided monosynaptically by glutamatergic RGC input and followed by a second peak reflecting center-induced activity from the recurrent connectivity of neighboring excitatory neurons. This corresponds experimentally to the magenta traces in Fig. 2C–D, where the distinction between the peaks is less pronounced in the electrophysiological recordings, appearing as compound postsynaptic currents. This second peak primarily depended on the strength of recurrent excitatory connections (*E-E*) and covaried with the strength of the *I-E* connection (Fig. 4B–C), which we refer to as feedback or recruited inhibition, as it requires the recruitment of inhibitory neurons by retinal input.

To investigate the role of the surround in modulating center responses, we activated the 3286 excitatory and 3286 inhibitory neurons (Fig. S7C). When center stimulation coincided with this surround input (cyan traces, Fig. 4B), the peak of the secondary response depended on the mutual coupling between excitatory and inhibitory neurons (*E-I* and *I-E*). The degree of secondary response attenuation varied based on the strength of these two connections (Fig. 4C). Surround inhibition played a critical role in effectively reducing the secondary response below a hypothetical threshold (dotted line in Fig. 4B). This attenuation was a result of fewer excitatory and inhibitory neurons responding above threshold to the center input, as result of both direct and indirect effects of surround inhibition.

To quantify the contribution of individual circuit motifs, we normalized the center response during surround network activation (Fig. 4C) by the peak when only center input was delivered. Any increase in the *I-E* connection strength increased secondary peak attenuation, whereas *E-I* connectivity did not have a strong effect on the attenuation of the secondary peak. The weight of recurrent inhibition (*I-I*), a prominent feature of the SCs layer (Fig. 3L), did not have an impact on surround suppression. Notably, strong recurrent excitation increased the attenuation of the secondary excitatory conductance peak during surround network activation (compare top and bottom rows and panels in Fig. 4B and 4C, respectively). In conclusion, these findings underscore that a robust network characterized by prominent recurrent *E-E* connectivity is essential for the reduction of center excitation when an active surround drives feedback inhibition (*I-E*).

## Discussion

Targeted optogenetic activation of center and surround zones in individual SCs neurons allowed us to test the presence and circuit implementation of surround modulation in the superficial layers of the superior colliculus. We demonstrate that reduction in center excitation occurs when the surround is activated by retinal input (Fig. 2), resulting from both direct and indirect mechanisms. Given our experimental setting, which involves midbrain slices that naturally disconnect other brain regions, this finding indicates that the observed changes in synaptic excitatory conductance—ranging from moderate to strong—are the result of local interactions occurring within the midbrain superior colliculus. This suggests that primary extraretinal circuits (i.e., SCs and V1) located within divergent subcortical and cortical pathways, could potentially perform surround suppression independently. In principle, this could enable a stimulus to undergo selective suppression in one pathway (e.g., thalamocortical) while being enhanced in another (e.g., retinotectal), though this would need to be confirmed *in vivo*.

As in cortical circuits (39), the SCs contains excitatory and inhibitory neurons that are spatially intermingled, a feature that complicates the unravelling of their function. By combining whole-cell measurements (Fig. 2) and connectivity analysis (Fig. 3) with a computational model (Fig. 4), we revealed two main local circuit motifs: recurrent excitation (*E-E*) and feedback inhibition (*I-E*) that contribute to surround suppression in distinct ways. *E-E* connections amplify responses in a specific retinotopic ‘center’ whereas *I-E* performs a dual function. It overlaps with center-specific *E-E*, thereby modulating the responses of the same center neurons subject to *E-E*, while simultaneously, reaching slightly beyond this zone, curbing the responsiveness of neighboring surround neurons outside the center.

The selective decrease in center excitation when the visual surround is active is consistent with models that involve amplification in recurrent networks (40), but inconsistent with models that are dominated by feedback inhibition (41). In the latter, transient increase in inhibition would account for reduction in overall excitation and inhibition levels – an outcome that our measurements did not support. We show that during surround network activation, recurrent interactions conspire to reduce the persistence of both excitatory and inhibitory inputs. Computational models of trial-by-trial variability suggest that, while both increases in inhibition or decreases in recurrent excitation can similarly affect the average output response suppression, the latter is more likely to minimize variability across trials (42).

Visual saliency computations in the SCs provide cortical circuits the flexibility to modulate SCs activity either by amplifying or suppressing with retinotopic correspondence (25). This could allow them to engage with SCs recurrent networks and modulate local center-surround dynamics in a top-down manner. For instance, by granting selective access for pyramidal neurons in V1 to interact with *E-E* connections, responses can be biased to specific incoming retinal inputs from certain areas of the visual field, or in contrast, they can be suppressed by activating *I-E* connections. Future studies employing the *excitation zone mapping* method presented here to investigate center-surround interactions within the V1-SCs pathway have the potential to reveal the intricate circuit logic underlying top-down cortico-collicular interactions, in concert with retinal-driven center-surround mechanisms. Understanding how phylogenetically older and neocortical circuits can cooperate or compete is important for comprehending visual saliency computations.

## Material and Methods

### Animal strains

Experimental procedures were approved by the Stockholm municipal committee for animal experiments and the Karolinska Institute in Sweden (N9179-2017 and N6747-2019 to A.A.K.), as well as, the Committee on Animal Research at the Universidad Miguel Hernández and carried out in compliance with the Research Council of Spain (CSIC), the Generalitat Valenciana (GVA) and European regulations (2022/VSC/PEA/0236 to A.A.K.). Both male and female vGAT-Cre (Jackson: Slc32a1tm2(cre)Lowl), vGluT2-Cre (Jackson: Slc17a6tm2(cre)Lowl) and wild-type mice (C57BL/6N; Charles River) were used throughout the experiments. Mice were kept on a 12-h day/night cycle under stable temperature (21 ± 1 °C) and air humidity (50–65%) with freely available food and water. The exact numbers of cells (*n*) and animals (*N*) used for each experiment are reported in the *Results* section and corresponding figure legend.

### Viral injections

The complete list of viruses we used is provided in Supplementary Table 1. Injections were performed in 2–5 months old wild-type mice and transgenic mice under isoflurane anesthesia and injected with buprenorphine post-surgery. Viral titers of adeno-associated viruses (AAV) and modified rabies virus were between 1 × 10^12^ and 1 × 10^13^ viral genomes per milliliter (vg/mL). To express ChR2 in the axons of RGCs in the superficial layer of the right hemisphere SC, we injected 1000 nL into the vitreous bodies of the left retinas using the following viral vectors:

1. rAAV2/hsyn-hChR2(H134R)-EYFP-WPRE-PA (UNC Vector Core),
2. rAAV2/hsyn-hChR2(H134R)-mCherry-WPRE-PA (UNC Vector Core),
3. rAAV2-EF1α-DIO-hChR2(H134R)-EYFP (UNC Vector Core), or
4. ssAAV-2/2-shortCAG-dlox-hChR2(H134R)_EYFP(rev)-dlox-WPRE-hGHp(A) (VVF, Zurich).

We used the vector rAAV2/hsyn-mCherry (UNC Vector Core) as in our control group for the intravitreal injections. To identify GABAergic neurons in the SCs, the vector rAAV5-CAG-FLEX-tdTomato (400 nL, UNC Vector Core) was injected into two collicular sites in order to maximize expression: 1) the rostral SCs (from bregma: anteroposterior (AP) -3.6 mm; right (R) -0.8 mm; dorsoventral (DV) -1.6 mm from the surface of the skull), and 2) the caudal SCs (AP -4.2 mm; R -1.8 mm; DV -1.5 mm) respectively on the same day.

For the rabies virus tracing, 50 nL of helper virus AAV5-EF1a-DIO-TVA-V5-t2A-RG was injected into the right SCs (AP -3.9 mm; R -0.8 mm; DV -1.05 to -1.15 mm from the surface of the brain) in vGAT-Cre or vGluT2-Cre mice, while taking great care to maintain viral spread to within the SCs layer. Three weeks later, 150 nL of the engineered rabies virus EnvA-coated G-deleted rabies virus with EGFP (EnvA-ΔG-Rb-EGFP) was injected into the same location.

For secondary injections, fiber optic implantations and electrophysiological recordings, two to three weeks were allowed to pass after the initial injection. Viral suspensions were delivered at 500 nL/min and 100 nL/min for intravitreal and brain injections, respectively. A puncture was made 2 mm posterior to the corneal limbus using a sterile 30-gauge (G) needle after pupil dilation was achieved with 0.5% Alcaine. After which 1000 nL vitreous was aspirated manually using a Hamilton microliter syringe. The glass capillary was then inserted through the same puncture by using a microliter syringe (1.5 mm volume) into the eye, and the tip was angled toward the vitreous humor and back of the eye, taking care to avoid any damage to the retina or lens. The capillary remained in the eye for 2 min in order to prevent reflux of the viral suspensions. Brain injections were performed by using a digital stereotaxic frame combined with a Quintessential Stereotaxic injector (Stoelting) via a glass capillary. The capillary firstly reached 0.2 mm deeper than the target location to give space for the viral suspensions, then was slowly retracted to the target location to complete the injections. Afterwards, the capillary was retracted by 0.2 mm again and remained in position for 8 min. During the injections, all the mice were kept on a heating pad at 37 °C with a subcutaneous injection of buprenorphine while maintaining the eyes were moisturized with Viscotears gel.

### Patch electrophysiology in acute midbrain slices

The animals were deeply anaesthetized by using a combination of pentobarbital injected intraperitoneally. To ensure rapid cooling of the midbrain, we perfused the animals transcardially with an ice-cold cutting solution, containing the following (in mM): NaCl 40, KCl 2.5, NaH_2_PO_4_ 1.25, NaHCO_3_ 26, glucose 20, sucrose 37.5, HEPES 20, NMDG 46.5, Na L-ascorbic acid 5, CaCl_2_ 0.5 and MgCl_2_ 5 (pH 7.3 with HCl). The brain was rapidly dissected out after decapitation. Coronal midbrain slices (400 μm) were prepared by using a vibratome (VT1000S, Leica, Germany) in the cutting solution and incubated in the same solution at 35 °C for 12 min. Slice orientation was chosen to profile responses along the medio-lateral axis corresponding to the spatial axis of the retina across the dorso-ventral direction of visual space, whereas the dorsal-ventral axis of the SCs slice corresponded to the various sublaminae of the SCs. The sections were subsequently maintained at room temperature in recording solution that contained the following (in mM): NaCl 124, KCl 2.5, NaH_2_PO_4_ 1.25, NaHCO_3_ 26, glucose 20, CaCl_2_ 2 and MgCl_2_ 1 (pH 7.3). Cutting and recording solutions were continuously infused with carbogen (95% O_2_ and 5% CO_2_) throughout the procedure.

Whole-cell patch clamp recordings were either performed on random neurons in the SCs of wild-type mice or tdTomato-expressing neurons of vGAT-Cre mice at room temperature using pCLAMP 11 (Molecular Devices, USA). Midbrain slices were continuously perfused with oxygenated recording solution at a rate of 2 mL/min during recording and visualized under an upright CleverScope motorized microscope (Micro Control Instruments, UK) equipped with a Dodt gradient contrast system (DGC-2), an oil condenser (U-AAC, Olympus, Japan) and two Chameleon3 cameras (Teledyne FLIR, USA). Whole-cell recordings were obtained under visual control using a 16× (0.80 NA) water dipping objective (CFI75 LWD 16X W, Nikon, Japan) combined with 2× c-mount extender (EX2C, Computar, USA) and c-mount adapters (U-TV1X-2 and U-CMAD3, Olympus, Japan) with Axon MultiClamp 700B amplifier and Axon Digidata 1320A digitizer (Molecular Devices, USA). Signals were sampled at 20 kHz. Pipettes used for recording were pulled from borosilicate glass capillaries (article no. 1403542, Hilgenberg, Germany) by a P-97 Micropipette Puller (Sutter Instrument, USA). Patch pipettes (5–8 MΩ) were filled with a potassium gluconate based intracellular solution contained (in mM): K D-gluconate 125, EGTA 1, KCl 10, HEPES 10, ATP-Mg 4, GTP-Na 0.3, phosphocreatine 10, CaCl_2_ 0.1 and QX314 3 (pH 7.2 with KOH; 290 mOsm). Due to the long recording times (usually one hour), the access resistance was evaluated every 10 min to ensure that initial values (only under ∼30 MΩ were considered) did not increase throughout the recording session.

ChR2-expressing axons of retinal ganglion cells were optically stimulated with 470 nm light from a high-power collimated LED source (Mightex, Canada) connected to a digital mirror device (Polygon1000, Mightex, Canada) by a 3 mm-core liquid lightguide (Newport, USA). The stimulation patterns were triggered via Master-8 pulse stimulator (A.M.P.I, Israel) and designed via the software PolyScan2 (Mightex, Canada). The tdTomato-expressing vGAT-positive neurons in the SCs were detected with 540 nm light from a high-power collimated LED source (Mightex, Canada) and an RFP filter set (Olympus, Japan). The 540 nm LED was connected to a 495 nm dichroic filter cube (Mightex, Canada) with the 470 nm LED during the transgenic animal experiments. Both LEDs were directed via a four channel BioLED light source control module (Mightex, Canada).

For membrane property measurements, a series of 1000 ms negative and positive injected current steps from -30 pA to +40 pA in 5 pA increments were delivered through the recording electrode under current-clamped conditions to each recorded neuron.

### Excitatory zone mapping

To accurately cover each neuron’s center zone and ensure a significant surround area, we created a sufficiently large field of view (FOV) that could accommodate their distinct morphologies. We achieved this using one-photon widefield optics with a low magnification objective (16×) and an in-house built dual-focus visual inspection system that allowed selective magnification and demagnification of the optical path (Fig. 2B). Demagnification (by 0.5×) captured a large FOV of > 1 mm², encompassing most of the SCs (magenta box in lower right of Fig. S1A), enabling patterned micro-optostimulation of the entire visual layer. The alternate path for magnification (by 2×) provided sufficient single-cell resolution to visually target neurons of interest for whole-cell recordings.

We combined whole-cell electrophysiology with a digital mirror device (DMD)-based dual-camera system to map the receptive fields of individual neurons. Optogenetic stimulation patterns were created and implemented during recordings by the DMD (Polygon1000, Mightex, Canada). The 16× objective, along with 2× front tube optics (Mightex, Canada) and a 0.5× single photo port tube lens camera adapter (Olympus, Japan), enabled a workable field-of-view of approximately 1.2 × 1 mm. Using this customized visual inspection system, whole-cell patch-clamp recordings were performed through the high magnification pathway while optostimulation patterns were executed via the low magnification pathway. All neurons were voltage-clamped to −65 mV and 0 mV to measure excitatory postsynaptic currents (EPSCs) and inhibitory postsynaptic currents (IPSCs), respectively, with an intermediate potential (−45 mV) applied for membrane potential recordings in most neurons when stable. Traces were not corrected (NC) for the offset created by the liquid junction potential, which was experimentally confirmed to be ∼10 mV.

For control experiments, whole-field stimulation patterns were delivered by 10 Hz trains of 8 stimuli (5 ms duration), followed by a single test stimulus at 5 s time interval, and each trace included four repetitive sweeps. For the center-surround experiments, SCs neurons were subjected to the following patterns: ‘GridScan’, ‘Center-Center’, ‘Surround-Surround’ and ‘Surround-Center’. Center (C) and Surround (S) areas were determined by using the GridScan protocol, which was delivered 10 Hz trains of 5 light pulses of 5 ms duration in 100 non-adjacent grids arranged in a 10 × 10 spatial configuration across the FOV (0.1 × 0.07 mm for each one). After the end of the GridScan, we promptly calculated the maximum EPSC and IPSC amplitudes for each grid after it was normalized and filtered using custom-written MATLAB (MathWorks). Excitatory postsynaptic current amplitudes arising from each grid location were thresholded on the basis of their evoked amplitude. Grid locations that yielded ≥ 10–20% of the peak EPSC amplitude for the recorded neuron, were considered to be part of the center zone, whereas locations that did not pass this threshold were considered to be part of the surround zone. To trigger center-surround interactions, the surround was optostimulated by a single pulse that was then followed by a two or five-pulse train stimulation delivered to the center. We experimented with a range of different surround stimulation durations (5, 20, 50 ms) and with a range of surround-center delays (0, 10, 20, 50 ms) to explore the parameter space. The inter-trial interval was 25 ± 5 s.

### Drugs used in electrophysiology

Drugs were bath applied and an overall duration of at least 5 min was allowed before testing their effects, which was followed by a washout phase of at least 10 minutes. Unless otherwise indicated, drugs were perfused during patch-clamp recordings under the given conditions using a two-gauge peristaltic pump (Pretech Instruments, Sweden). The complete list of drugs used is provided in Supplementary Table 2.

### Estimation of synaptic conductances

To deconvolve inhibitory and excitatory synaptic conductance, recordings were performed during ‘center-center’, ‘surround-surround’, ‘surround-center’, and ‘global’ optostimulation. A QX314-based intracellular solution was used in the recording pipette to improve intrinsic conductance isolation and block action potential discharge conditions (Fig. 2 and Fig. S1–2). Evoked synaptic currents used in this analysis were obtained by averaging values from 5 to 10 sweeps for each recorded cell. Due to the long recording times (usually one hour), the access resistance was evaluated every 10 min to ensure that initial values (only < ∼30 MΩ were considered) did not increase throughout the recording session at each membrane potential. Customized MATLAB scripts were written to estimate the total, excitatory and inhibitory synaptic conductances based on the equations used in (34, 35). It should be noted that the postsynaptic currents were adjusted for the liquid junction potential (∼10 mV) and taken into consideration when estimating for their underlying evoked synaptic conductance.

### Neuronal reconstruction

To identify the dendritic fields from the recorded neurons, we used 0.3% Neurobiotin (Vector Laboratories, no. VESP-1120-50) in the intracellular solution during whole-cell recordings. We post fixed the midbrain slices and then used PBS to wash them. To visualize the neurons, we stained them with streptavidin Cy5 (1:400, Jackson ImmunoResearch, no. 016-170-084) and imaged them using confocal microscopy with a 20× objective (LSM-800, Zeiss, Germany). The images were processed with ImageJ software, and the neurons were reconstructed by using the plugin Simple Neurite tracer. The dendritic areas of each labeled neuron were calculated by tracing a convex polygon around the outermost tips of the dendritic field. The SC cell types were then classified on the basis of their geometry (36).

### Fiber optic implantation and *cFos* activation

Two weeks after the intravitreal injection, a 200 μm diameter fiber optic (Thorlabs, FG200UEA) was implanted into the right SCs (from bregma: AP -3.6 mm; R -1.0 mm; DV -1.0 mm from the brain surface). The mice recovered for at least one week before the *cFos* induction experiment. The mice were habituated to handling in an open field area (60 cm × 60 cm × 60 cm; W×D×H) and the connected optic fiber for at least 3 days before proceeding with the experiment. To minimize stress, bedding from the cage was placed in the box. Optogenetic stimulation was not initiated for 90 min after the mice were placed in the dark box to avoid any prior visual effects on the observed *cFos* expression. The RGC axonal terminals in the SCs were stimulated with a train of 1 s square pulses of 40 Hz frequency (10 ms ON and 15 ms OFF) every 10 sec for a total of 40 min. After the stimulation period, the mice were left undisturbed for 60–70 min to ensure sufficient *cFos* protein expression. A maximum of 90 min after the end of the stimulation period was allowed before the mice were euthanized with pentobarbital and perfused with ice-cold 4% paraformaldehyde (PFA) in 0.01 M (1×) phosphate-buffered saline (PBS) buffer. The brain and retinas were subsequently dissected, and post fixed in 4% PFA overnight at 4°C.

### Immunofluorescent staining and *in situ* hybridization

Mice were anesthetized by pentobarbital and perfused with 1× PBS and 4% PFA. The brains (and eyes, to confirm pan-retina transfection) were harvested and fixed in PFA at 4 °C overnight, then washed with PBS for 3 times. The brain was placed in 15% sucrose solution for 24 h, then transferred into 30% sucrose solution for 24–72 h. The brains were subsequently embedded in optimal cutting temperature compound (OCT), frozen with dry ice and kept at -20 °C for short-term storage. The cryostat sectioning was performed on a Epredia NX70 cryostat. The brains, between the thalamus and brainstem, were sectioned in 14 μm thickness, about 80% (12 to 15) sections in the SCs were collected on glass slides, and every 6^th^ section was sampled for the following staining (the interval is about 0.105 mm). The other parts of the brain were sectioned in 40 μm thickness and stored in 1× PBS with 0.01% sodium azide.

The fluorescence *in situ* hybridization was performed with RNAscope Multiplex Fluorescent Reagent Kit v2 (ACD, Cat. No. 323100). The probes for *vGluT2* (Mm-Slc17a6-C2, ACD, Cat. No. 319171-C2) and *vGAT* (Mm-Slc32a1, ACD, Cat. No. 319191) were used to identify the cell types. The manufacturer’s protocol was followed. Co-staining of *vGAT* and *vGluT2 in situ* hybridizations was also performed for control experiment, and the results were shown in Fig. S5.

Immediately following the *in situ* hybridization, the brain sections were incubated in chicken anti-V5 primary antibody (Abcam, ab9113) in 1:750 at room temperature overnight, then washed with 1× TBST (0.3% TritonX-100), and incubated in donkey anti-chicken Cy3 secondary antibody (Jackson ImmunoResearch, no. 703-165-155) in 1:1000. All the antibodies were diluted with 1× TBST (0.3% TritonX-100, 3% BSA, 0.01% sodium azide). After PBS washing and DAPI staining, the sections were mounted with an antifade mounting medium (VWR, BSBT AR1109). For the *cFos* staining, rat or rabbit anti-*cFos* primary antibodies (Synaptic Systems, no. 226-017 or no. 226-003, 1:500) and GABA primary antibody (Sigma-Aldrich, A2052, 1:2000) were used in 1× PBST (0.3% TritonX-100, 5% donkey serum, 0.01% sodium azide) at room temperature overnight, then the sections were stained with Cy3 donkey anti-rabbit (Jackson ImmunoResearch, no. 711-165-152) or AF647 donkey anti-rat (Jackson ImmunoResearch, no. 712-605-153). The same following steps are previously described.

The retinas were first removed from the eye in the perfused animals and then flattened with the help of four incisions, and finally washed in 1× PBS for 3 times. For the retina immunostaining, the retina was first treated with pre-heated (about 70 °C) 1× antigen retrieval citrate buffer (Sigma, C9999), then followed the same protocol as with the brain section described above. Rabbit anti-S-opsin primary antibody (Sigma, AB5407) and donkey anti-rabbit Cy3 secondary antibody were both used in 1:1000 to visualize the short-wave photoreceptors, which reveals retinal orientation.

### Network model

We used an integrate and fire model to simulate the dynamics of neurons. The subthreshold membrane potential (*V*_*m*_) of each neuron was described by the following differential equation:

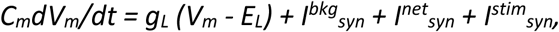

where *C*_*m*_ is the membrane capacitance, *g*_*L*_ is the leak conductance, 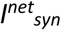 is the total synaptic input from neurons within the network, 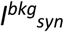 is the background input from other networks, 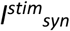 is input corresponding to center or surround inputs (*see* Supplementary Table S3). When the membrane potential reached *V*_*th*_, spike was elicited and the *V*_*m*_ was reset to *E*_*L*_ for *t*_*ref*_ millisecond. In the network all neurons were identical. Neuron parameters are provided in Table S4.

Synaptic inputs were modelled as conductance transients with following dynamics:

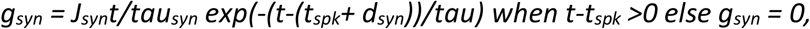

where *t*_*spk*_ is the spike time, *d*_*syn*_ is the synaptic delay, *J*_*syn*_ *{syn: ee, ei, ie, ii, ex, ix)* is the amplitude of the postsynaptic potential (PSP), and *tau*_*syn*_ is the synaptic time constant. In our model all synapses of a particular type (external, excitatory, or inhibitory) have the same parameters (*see* Supplementary Table S5). In our model we fixed the amplitude (*J*_*syn*_) of the external excitatory (*J*_*ex*_), inhibitory (*J*_*ix*_) synapses and *J*_*ii*_ (inh. → inh.). The value of *J*_*ee*_ (exc. → exc.), *J*_*ei*_ (exc. → inh.), and *J*_*ie*_ (inh. → exc.) were systematically varied. Synapse parameters are provided in Table S5.

We considered a network with 6400 excitatory and 6400 inhibitory neurons. Neurons were placed on an 80 × 80 grid. Because we have the same population size for the two neuron types, both types of neurons were placed on the same size grid. Neurons were connected based on their spatial distance. To avoid boundary effects, the 80 × 80 grid was folded as a torus. The connection probability decreased as a function of distance according to a Gaussian function. The standard deviation of the Gaussian was set to 16 and 20 for excitatory and inhibitory synapses, respectively. This was based on our experimental data which showed that inhibitory neurons have a slightly broader extent of their spatial connectivity than that of excitatory synapses (Fig. 3 and Fig. S5). 16 and 20 grid points constitute 2% and 2.5% of our network space. The area of the Gaussian function (*i*.*e*. the total number of connections sent out by a neuron) was decided based on the input convergence we measured in our experiments (Fig. 3H–I). Network connectivity parameters are provided in Table S6.

Each neuron received Poisson type excitatory and inhibitory spiking inputs to mimic inputs from outside the network and to obtain a low firing rate (< 1 Hz) background activity. *See* Supplementary Table S7 for parameters.

To model a short flash like input to the center region, we selected 314 excitatory and 314 inhibitory neurons, located in a circle (radius = 10 grid points) at the center of our network model. These neurons received a single synchronous spike from 10 presynaptic neurons (from the retina). Each spike elicited 1.1 mV depolarization. Each neuron received such a synchronous spike event at the stimulus onset (*Tcenter*). Each individual neuron received the synchronous spike event at a time *Tcenter + r*, where *r* is a random number drawn from a uniform distribution (U[0,1]). Center stimulus was presented to the neurons in three conditions: center only, center stimulus just at the end of the surround stimulus (*see* below) and center while the surround stimulus was still on. *See* Supplementary Table S3 for more details.

To mimic the surround stimulus, we selected 3286 E-neurons and 3286 I-neurons from a square region in the network with a side of 60 grid points. The center neurons (*see* above) were located at the center of this square. For the surround stimulus we excluded the center neurons. The surround stimulus was injected in the selected neurons in the form of uncorrelated Poisson type spike trains. *See* Supplementary Table S3 for more details.

The network model was simulated using the simulator NEST (43).

To quantify these network interactions, we used a spiking neuron model with conductance-based synapses, which were connected to each other based on physical distance according to a Gaussian connectivity kernel (*see* Table S3). Because we were interested in the contribution of different connections, we kept the connection probability and distance-dependent connectivity kernel fixed and systematically varied the connection strengths (*see* Table S3 for stimulus input; Table S4 for neuronal parameters; Table S5 for synaptic parameters; Table S6 for network connectivity; and Table S7 for external input parameters). Consistent with experimental findings (44), the network remained in an inhibition-dominated activity regime in which inhibitory neurons displayed a higher background activity than excitatory neurons for all synaptic strengths.

### Statistical analysis and data plotting

For each mouse, 10–16 sections were stained and visualized. Confocal images (20×) were used for the manual scoring of co-expression of vGAT/vGluT2, V5 and RV-GFP. Whole-brain images (10×) were acquired with a fluorescent microscope (Leica DM6000B) and a digital camera (Hamamatsu Orca-FLASH 4.0 C11440). They were manually registered to a bregma AP coordinate (from bregma -3.2 to -4.9), and subsequently aligned to the Allen Reference Atlas with the WholeBrain R package (45, 46). Every detected Rb-EGFP positive neuron was registered, and the coordinates were used for the following analysis. As the main source of monosynaptic input and local input, only the RV-GFP positive cells in the SCs were included in the calculation. For the comparison between mice, the number of input neurons were normalized to the total number of detected Rb-EGFP positive neurons in SCs. All the density estimations were done using a Gaussian kernel and were then plotted in R.

To minimize the effects of M-L differences of the injection sites between animals, the centroid of the starter neurons was first computed using k-means clustering analysis, which then allowed us to plot the relative M-L coordinate for each neuron. As for the depth estimation, the coordinates of the SC surface were first obtained with the WholeBrain R package, then the depth of each neuron was calculated after subtracting the difference. In both, M-L and depth density plots, only 3 central brain sections with the highest starter density were included according to the AP density plot.

After confocal imaging the *cFos* fluorescence was quantified by a 3d object counter plug-in of Fiji. Colocalized *cFos*+ and GABA+ cells were manually counted in under the optic fiber implant (Fig. 2F–G and Fig. S6). The statistical analysis and visualization were done in Python (code available upon request).

We used three model stimulation protocols: center only (center), center stimulus just at the end of the surround stimulus (center after surround) and center while the surround stimulus was still on (center with surround). Each paradigm was repeated 20 times. For each stimulus presentation, we recorded excitatory conductance from a single neuron. We estimated the across trial mean and variance of the conductance input to the recorded neuron. Finally, the results were rendered as the mean and variance of the excitatory conductance at specific time points after the onset of the stimulus in all three stimulus paradigms. In addition, we recorded spiking activity from all the neurons and estimated the firing rate of neurons. The network model was simulated using the simulator NEST. All differential equations were integrated using Runga-Kutta method with a time step of 0.1 ms.

Electrophysiology data analyses were performed using Clampfit 11 (Molecular Devices, USA), GraphPad Prism (GraphPad Software 7.00, USA), ImageJ (1.53c, NIH, USA), RStudio (R 3.6.1) or MATLAB (R2018b, MathWorks, USA). Data were all presented as mean values ± s.e.m. Before performing paired *t*-test or one-way ANOVA analysis, the normality and the homogeneity of variance were first evaluated using Kolmogorov–Smirnov’s test and Brown– Forsythe’s test. If the *p* value of Kolmogorov–Smirnov’s test was less than 0.05, Wilcoxon’s signed-rank test would be used instead of paired *t*-test. If *p* value of Brown–Forsythe’s test was less than 0.05, Friedman test and Kruskal–Wallis test would be used instead of a repeated-measures one-way ANOVA (RM one-way ANOVA) and ordinary one-way ANOVA, respectively. *p* values were corrected for deviation using the Geisser–Greenhouse correction.

Once the *p* value of a one-way ANOVA analysis was less than 0.05, Tukey’s multiple comparisons test would be performed. *F* and df (df1, df2) are degrees of freedom for one-way ANOVA and paired *t*-test, respectively. *p* values less than 0.05 were considered significant with asterisks in figures denoting as follows: \**p* < 0.05, \*\**p* < 0.01, \*\*\**P* < 0.001, \*\*\*\**p* < 0.0001, *ns*, no significance. Traces showing electrophysiological recordings plotted in the figures were not corrected for the liquid junction potential, which was experimentally found to be -9.8 mV (∼-10 mV). It was taken, however, into consideration when estimating for their underlying evoked synaptic conductance.

## Acknowledgements

We thank Drs. Michael Haüsser, Abdel El Manira and Julien Bouvier for comments on the manuscript. We thank Dr. Sten Grillner for his support throughout this study. We thank Drs. Mats Hellström and Sias Jordaan for technical assistance.

## Funding

Swedish Research Council Starting Grant in Medicine no. 2016-03134 (AAK)

Swedish Research Council Project Grant in Medicine no. 2020-02927 (AAK)

CIDEGENT funding from the Valencian Community in Spain no. 2021-028 (AAK)

Strategic research area neuroscience (StratNeuro) Start-up program 2017 (AAK)

Swedish Brain Foundation Hjärnfonden no. FO2018-0254 (AAK)

Hedlunds Foundation no. M-2017-0691 (AAK)

Olle Engvist Foundation no. 196-0148 (AAK)

Stiftelsen Kronprinsessan Margeratas Arbetnämnd för synskadade no. 2019-027 and 2020-182 (AAK).

## Author Contributions

PC performed electrophysiological recordings, viral injections, optogenetic experiments and data analysis. KS performed neuroanatomical experiments, RNAscope *in situ* hybridization, viral injections, fiber implantations and data analysis. DMM performed *in vivo cFos* induction experiments, visualization of neuroanatomy, and data analysis. AG. performed electrophysiological recordings. IL and TF contributed to neuroanatomy and *in vivo* experiments. AAK performed the conductance analysis. IL, KM, TF, and AAK contributed to the conceptualization and experimental study design. KM contributed vectors and resources for visualization of neuroanatomical data. AK contributed to the computational study design and performed the numerical experiments. AAK designed the figures, supervised and funded the study. AAK wrote the manuscript with help from AK. All authors reviewed and commented on the manuscript.

## Data and code availability

Data reported in this paper and all original code are available and will be shared by the lead contact upon request.

## Supplementary Materials

Please see Supplementary document for supplementary figures and tables.

