## Supplemental for "Recurrent circuits encode visual center-surround computations in the mouse superior colliculus"

### Supplemental material: Figures S1- S7 and Tables S1-S7

**Fig. S1.**

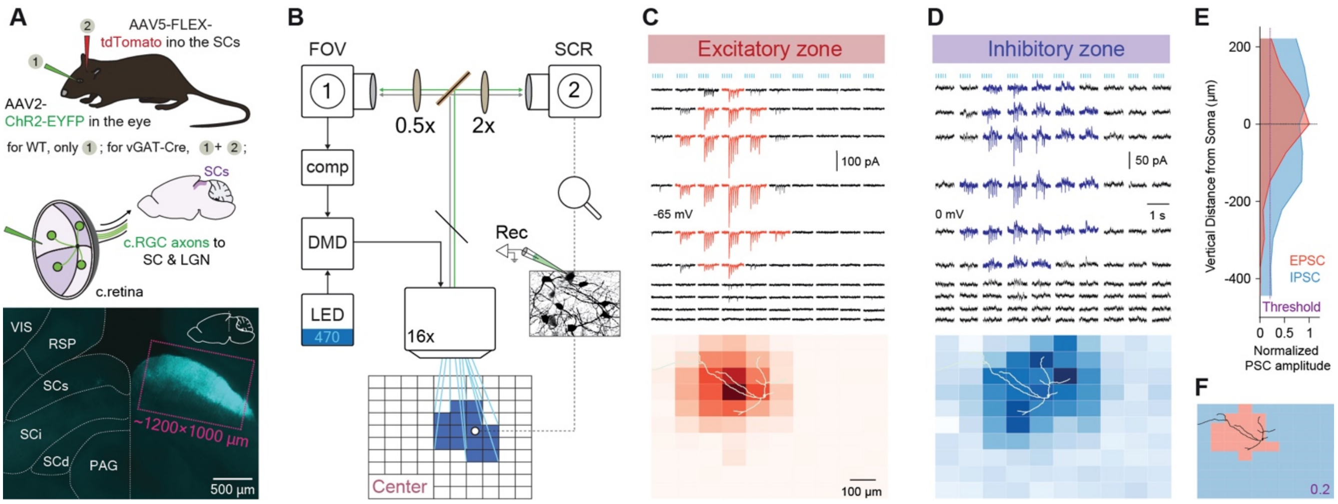

**Fig. S1. Mapping excitatory and inhibitory zones of SCs neurons**

*(A, Top)* Procedure showing expression of ChR2 in RGCs using an intravitreal viral injection of the vector rAAV2-hsyn-hChR2(H134R)-EYFP (in WT mice, *N* = 14) and unilateral labeling of contralateral SCs inhibitory neurons with the vector rAAV5-FLEX-tdTomato (in this case vGAT-Cre mice were used, *N* = 5). *(Bottom)* Coronal midbrain section with ChR2-expressing RGC axonal terminals seen occupying the SCs.

*(B)* Dual magnification setup for monosynaptic excitation zone mapping of single cells. This enabled the simultaneous visualization of a large field of view (FOV) encompassing the SCs (magenta rectangle in *A*), and single cell resolution (SCR) for identifying center excitation zones using whole-cell recordings and patterned optostimulation via a digital mirror device (DMD). *See* Methods for more information.

*(C–D)* A wide-field (WF) SCs neuron responding with excitatory and inhibitory postsynaptic currents (EPSCs measured at -65 mV shown in (A); IPSCs measured at 0 mV shown in (B) in response to a 20 Hz 5ms-light-pulse train delivered pseudo-randomly as a 120 × 100 µm grid optostimulation. Bottom panels: Response maps superimposed with the reconstructed morphology of the intracellularly stained WF neuron with Neurobiotin during whole-cell recording. Heat maps with darker shades coding for stronger responses in amplitudes.

*(E)* Distance from soma as a function of normalized postsynaptic amplitude in this WF SCs neuron. Red for excitatory; Blue for inhibitory. Vertical dashed line indicates threshold applied at a normalized value of 0.2.

*(F)* Center and surround zone of the WF zone determined by following thresholding approach applied to the excitatory PSCs shown in (C). Areas occupying a larger value compared to threshold are considered to be part of the center zone. *Abbrev:* c., contralateral; comp, computer; Rec, recording; WT, wild-type; ChR2, channelrhodopsin-2; RGC, retinal ganglion cell; VIS, primary visual cortex; RSP, retrosplenial cortex; SCs, superior colliculus superficial layer; SCi, superior colliculus intermediate layer; SCd, superior colliculus deep layer; PAG, periaqueductal gray matter; LGN, lateral geniculate nucleus.

**Fig. S2.**

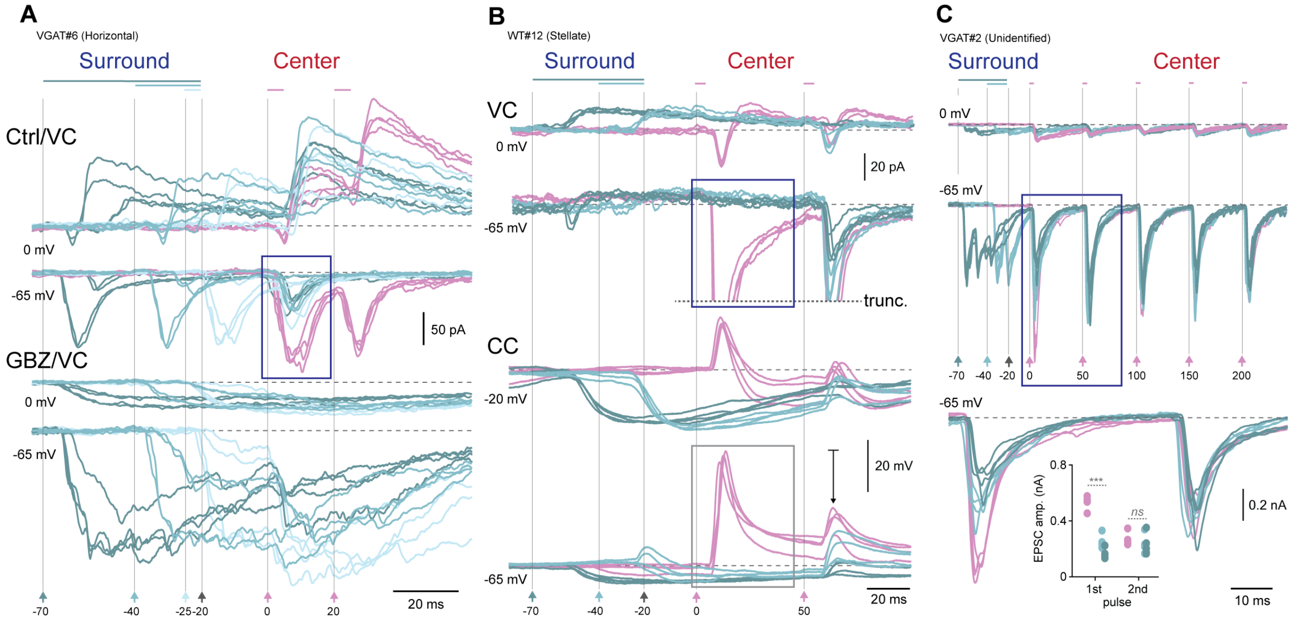

**Fig. S2. Three different examples of synaptic modulation between center and surround zones.**

*(A, Top)* Individual current traces held at reverse potentials for chloride-mediated GABAergic inhibition (-65 mV) and glutamate-mediated ionotropic excitation (0 mV) that span continuously surround and center optostimulation of the neuron (2 pulses separated by 20 ms) shown in Fig. 2C–D. Here, surround network activation can be seen to effectively reduce the center EPSC amplitudes (blue rectangle) when comparing to center only responses (in magenta). Evoked responses to surround stimulation of all durations (5, 20 and 50 ms; shades of green) are also seen to drive bistable excitatory and inhibitory responses. *(Bottom)* Voltage-clamp (VC) recordings in response to surround optostimulation after bath application of 10 µM Gabazine (GBZ), thereby removing the effect of GABAergic inhibition. Same color-coding scheme as in Fig. 2. Time stamps are centered around the onset of the first center pulse.
*(B, Top)* Synaptic current recordings performed in voltage-clamp of a stellate neuron that is subject to the 20 Hz optostimulation (2 pulses). Center EPSCs shown in the rectangle are truncated (dotted line). *(Bottom)* Same neuron recorded in current-clamp (CC) reveals the shunting effect that surround activation has on center responses (gray rectangle) when recording at -65 mV and at a depolarized membrane potential of -20 mV. This is attributed to chloride current triggered by the surround (revealed when held at 0 mV in VC; top traces).
*(C)* Current traces from a vGAT+ SCs neuron in response to 20 Hz optostimulation (5 center pulses) that shows signs of surround suppression without the direct effect of synaptic inhibition. Top and middle traces: Current recordings performed at -65 mV and 0 mV reveals that this neuron does not receive direct inhibition from the surround yet its center response is attenuated suggesting that it receives less excitatory input. Bottom: Enlarged view of current traces found in blue rectangle in middle panel focusing on first and second paired pulse center stimulations. Surround stimulation has a significant effect on the first center pulse by reducing its amplitude by 50% (***, $p < 0.0001$; *t*-test). Note that liquid junction potential correction was not performed, which was experimentally measured to be ~10 mV.

**Fig. S3.**

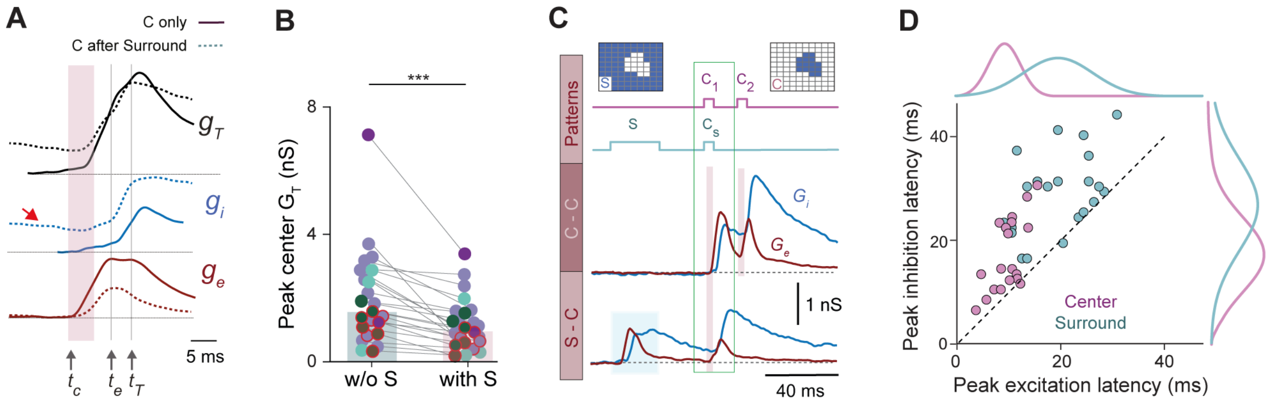

**Fig. S3. Center-surround dynamics are defined by transient excitatory and inhibitory conductances.**

*(A)* Total synaptic conductance ($G_T$, black line) triggered by center only ($C_1$) and surround-center ($C_s$) conditions is decomposed into excitatory ($G_e$, red) and inhibitory ($G_i$, blue) conductances. Conductances belong to recording shown in Fig. 3A, which are estimated by Wehr and Zador (2003) by using current recordings performed at -65 mV and 0 mV that were corrected for the liquid junction potential. Time stamps $t_c$, $t_e$ and $t_T$ symbolize the time onset for center stimulation, the time instant when peak excitation and maximal total conductance is reached, respectively. Solid lines are with center only condition and dashed lines are when center follows surround optostimulation. Red arrow indicates non-zero inhibitory conductance evoked by surround network activation at the time of $t_c$.

*(B)* Comparison of total synaptic conductance $G_T$ between $C_1$ and $C_s$ conditions. Corresponding excitatory and inhibitory conductances at peak $G_T$ are shown in Fig. 2F–G. Color coding corresponds to morphologically or genetically identified SCs neuron cell-types (*see* Fig. 2F).

*(C)* Full time course of excitatory and inhibitory conductance during both stimulation conditions, $C_1$ and $C_s$. *Legends:* C-C (paired pulse center optostimulation) and S-C refers to surround followed by center optostimulation after a 20 ms refractory period.

*(D)* Latency to peak inhibition as a function of latency to peak excitation. A relative delay $\Delta t$ of 20 ms is observed when comparing the center (magenta) and surround (cyan) conditions. This suggests that the signals originating from the surround region interact with the center responses with this time difference. Curves are a gaussian probability distribution function fitted to the data points.

**Fig. S4.**

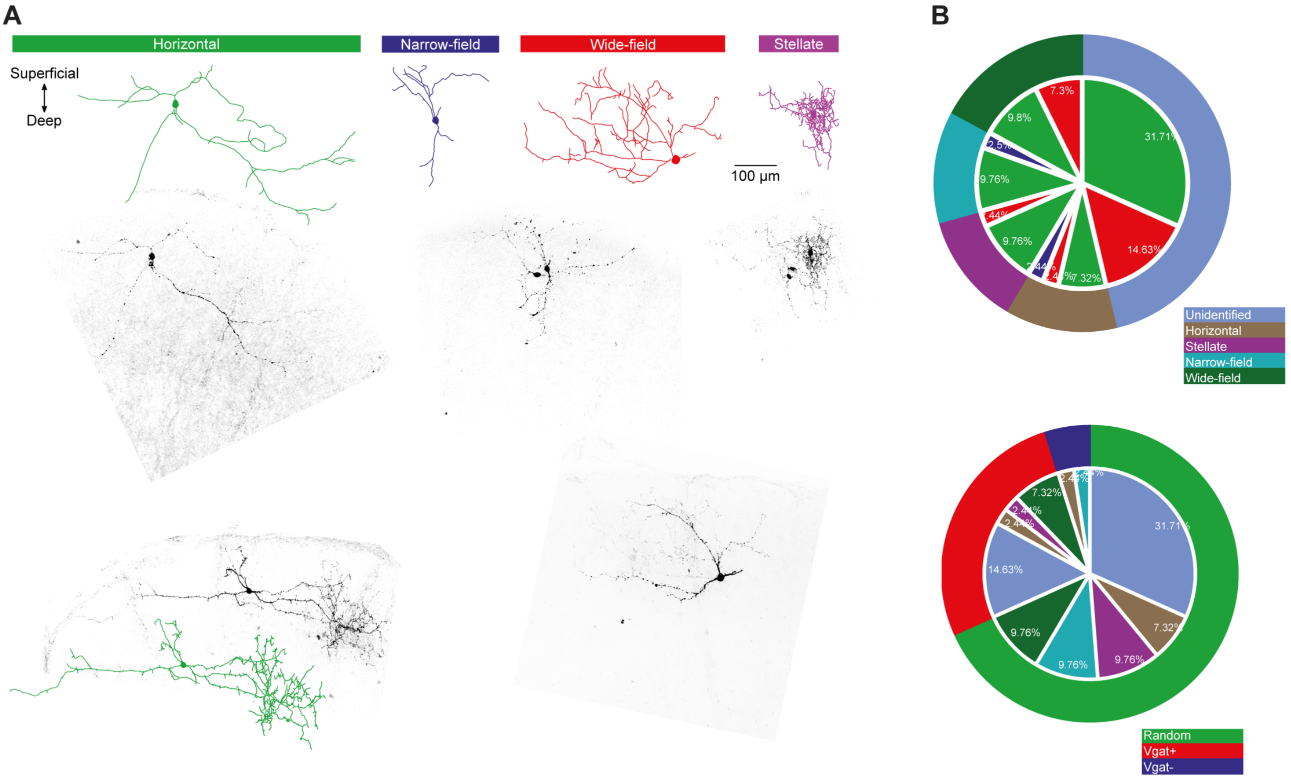

C

| Morphol-ogy | | 100.00% | Genotype | 41 | 500.00% | |
|---|---|---|---|---|---|---|
| | 41 | 41 | | 41 | 41 | 100.00% |
| Un-identified | 19 | 46.34% | Random | 13 | 68.42% | 31.71% |
| | | | Vgat+ | 6 | 31.58% | 14.63% |
| | | | Vgat- | 0 | 0.00% | 0.00% |
| Horizontal | 5 | 12.20% | Random | 3 | 60.00% | 7.32% |
| | | | Vgat+ | 1 | 20.00% | 2.44% |
| | | | Vgat- | 1 | 20.00% | 2.44% |
| Stellate | 5 | 12.20% | Random | 4 | 80.00% | 9.76% |
| | | | Vgat+ | 1 | 20.00% | 2.44% |
| | | | Vgat- | 0 | 0.00% | 0.00% |
| Narrow-field | 5 | 12.20% | Random | 4 | 80.00% | 9.76% |
| | | | Vgat+ | 0 | 0.00% | 0.00% |
| | | | Vgat- | 1 | 20.00% | 2.44% |
| Wide-field | 7 | 17.07% | Random | 4 | 57.14% | 9.76% |
| | | | Vgat+ | 3 | 42.86% | 7.32% |
| | | | Vgat- | 0 | 0.00% | 0.00% |
| Genotype | | 100.00% | Morphology | 41 | 300.00% | |
| | 41 | 41 | | 41 | 41 | 100.00% |
| Random | 28 | 68.29% | Unidentified | 13 | 46.43% | 31.71% |
| | | | Horizontal | 3 | 10.71% | 7.32% |
| | | | Stellate | 4 | 14.29% | 9.76% |
| | | | Narrow-field | 4 | 14.29% | 9.76% |
| | | | Wide-field | 4 | 14.29% | 9.76% |
| Vgat+ | 11 | 26.83% | Unidentified | 6 | 54.55% | 14.63% |
| | | | Horizontal | 1 | 9.09% | 2.44% |
| | | | Stellate | 1 | 9.09% | 2.44% |
| | | | Narrow-field | 0 | 0.00% | 0.00% |
| | | | Wide-field | 3 | 27.27% | 7.32% |
| Vgat- | 2 | 4.88% | Unidentified | 0 | 0.00% | 0.00% |
| | | | Horizontal | 1 | 50.00% | 2.44% |
| | | | Stellate | 0 | 0.00% | 0.00% |
| | | | Narrow-field | 1 | 50.00% | 2.44% |
| | | | Wide-field | 0 | 0.00% | 0.00% |

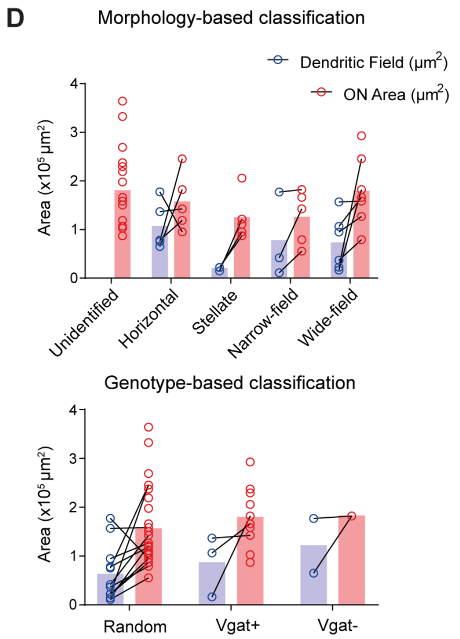

**Fig. S4. Identification of SCs cell types used for electrophysiology analysis**

*(A)* Reconstructed morphologies from various subtypes of SCs neurons that were recorded and presented in Fig. 3, including wide-field, narrow-field, stellate and horizontal neurons, that were stained with Neurobiotin.

*(B–C)* Quantification and classification of cell types according to morphology (i.e. horizontal, stellate, etc.) and genotype (i.e. vGAT+/- or random type). A total of *n* = 41 were performed of which a subset of *n* = 24 that maintained stability (stable access resistance over the period of the whole-cell recording

that usually lasted 60 min or 120 min in the case that drugs were bath applied) were included in the analysis shown in Fig. 2.

*(D)* Relationship between dendritic field and the neuron's corresponding retinorecipient center zone. Individual center zones typically exceed the area of the dendritic zone. Note that this correlation was performed for recorded neurons where morphological data were successfully obtained.

**Fig. S5.**

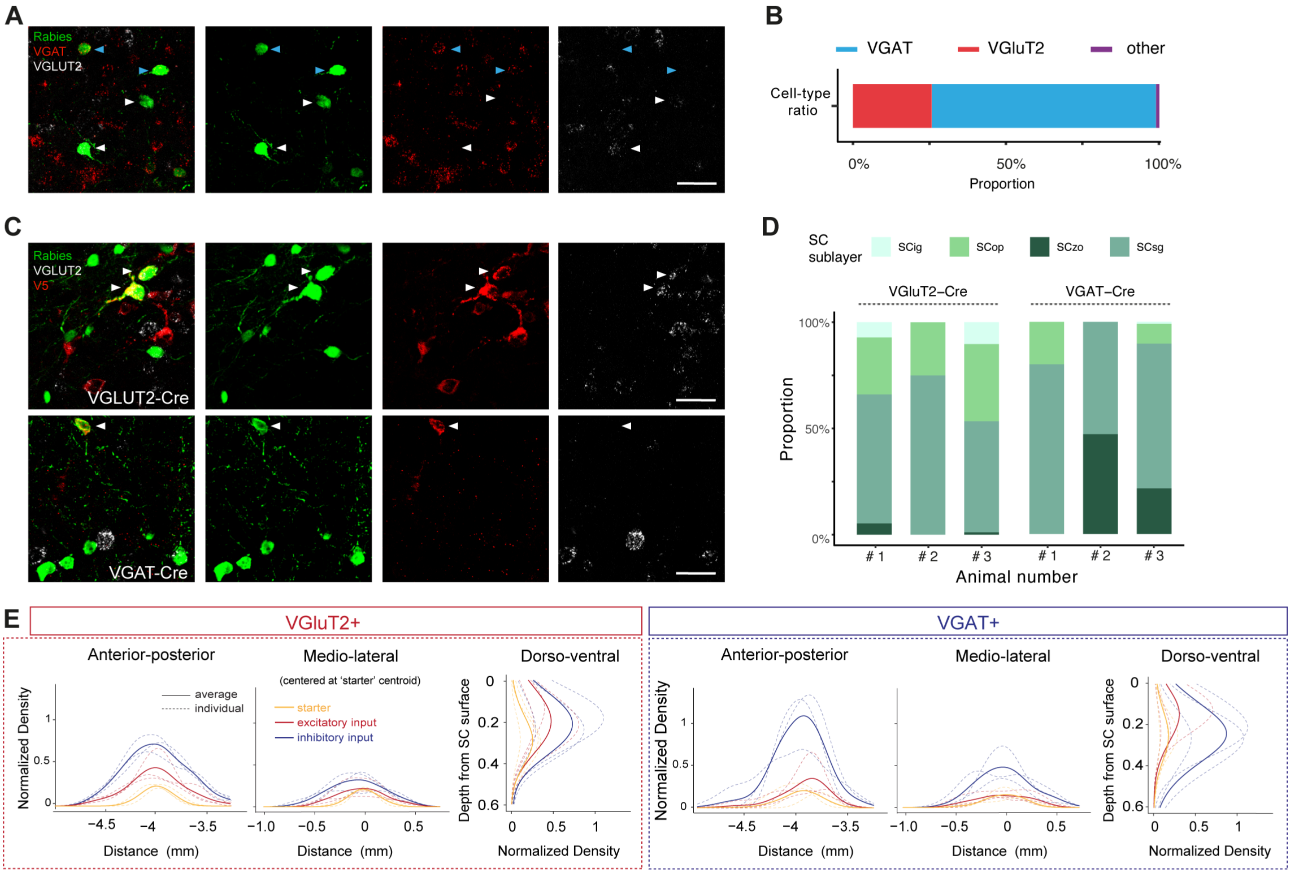

**Fig. S5. Classification of vGluT2 and vGAT cell types and their distribution across the SCs.**
*(A)* Co-expression of *vGAT* and *vGluT2 in situ* hybridization in the SCs using RNAscope *in situ* hybridization performed to reveal the identity of input neurons. White arrows indicate Rb-EGFP+ cells that are positive for *vGluT2* and negative for *vGAT*, suggesting that these input neurons are excitatory. By contrast blue arrows show Rb-EGFP+ cells that are inhibitory since they are positive for *vGAT* and negative for *vGluT2*. Scale bar: 25 µm.
*(B)* The proportion of vGluT2+ (red, *n* = 70) and vGAT+ (blue, *n* = 199) input cells that colocalize with Rb-EGFP. Unidentified neurons shown in purple (*n* = 3/272). *N* = 1 animal for vGAT and *N* = 1 for vGluT2, 3 sections were used per animal. Total number of counted neurons is: 272.
*(C)* RNAscope *in situ* hybridization for identity confirmation of starter neurons (*see* white arrows). Top row: Colocalization of vGlut2 detected in cells that express Rb-EGFP and the Cre-inducible V5 tag expressed in vGluT2-Cre mice, confirm starter neurons are excitatory. Bottom row: The same approach

applied in a vGAT-Cre. Colocalization Rb-EGFP+ with V5+ but negative for vGluT2, suggests that the starter neuron is putatively inhibitory. Scale bar: 25 µm.

*(D)* the distribution of starter neurons in each animal, shown as proportion in different sublayers.

*(E)* Normalized density plot of starter and input neurons along anterior-posterior axis (AP; Left), medio-lateral axis (ML; Center) or depth from SC surface (Right). Distribution of excitatory and inhibitory neurons are shown in the top and bottom rows, respectively. The ML density plot is computed on the relative coordinates to the k-means centroid of starter cells. Dashed lines represent individual animals, and solid lines show the average.

**Fig. S6.**

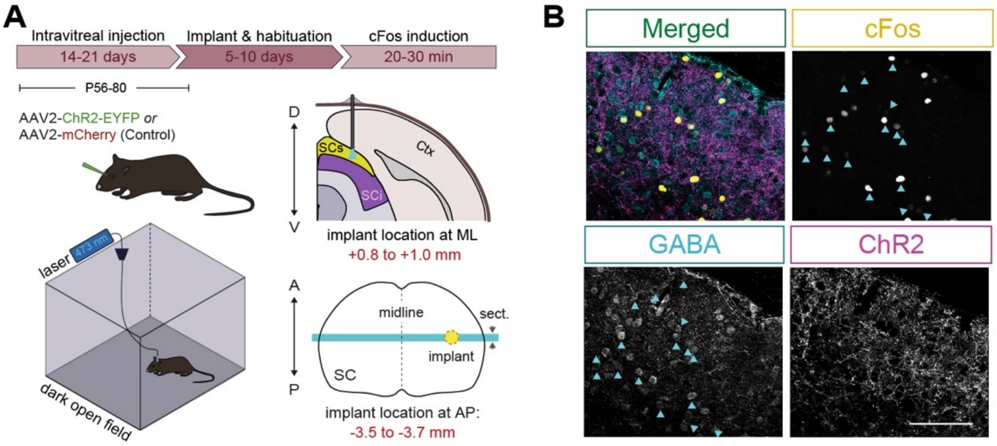

**Fig. S6. Functional recruitment of excitatory and inhibitory SCs neurons by RGCs (supplemental to Fig. 3)**

*(A)* Pipeline that we used to measure *in vivo cFos* expression induced by local optostimulation of RGC axonal terminals in the SCs. Top left: As in Fig. 1a, expression of ChR2 in RGCs was achieved using an intravitreal viral injection of the vector rAAV2-hsyn-hChR2(H134R)-EYFP (in WT mice, *N* = 5) or rAAV2-hsyn-mCherry for control (in WT mice, *N* = 3). Bottom left: schematic illustrating assay performed in a dark box. Right panels: cartoon of the optic fiber placement site shown in coronal (top) and horizontal (bottom) midbrain sections.

*(B) In vivo cFos* expression resulting from optostimulaon of RGC axonal terminals under the tip of the optic fiber implanted in the SCs (*see* Fig. 3E-H). From top left: a) merged image, b) *cFos* staining, c) GABA immunostaining (arrows show colocalization), and d) ChR2-expressing retinal ganglion axonal terminals. ChR2 shown in magenta; *cFos* in yellow; GABA in cyan. Scale bar: 50 µm.

**Fig. S7.**

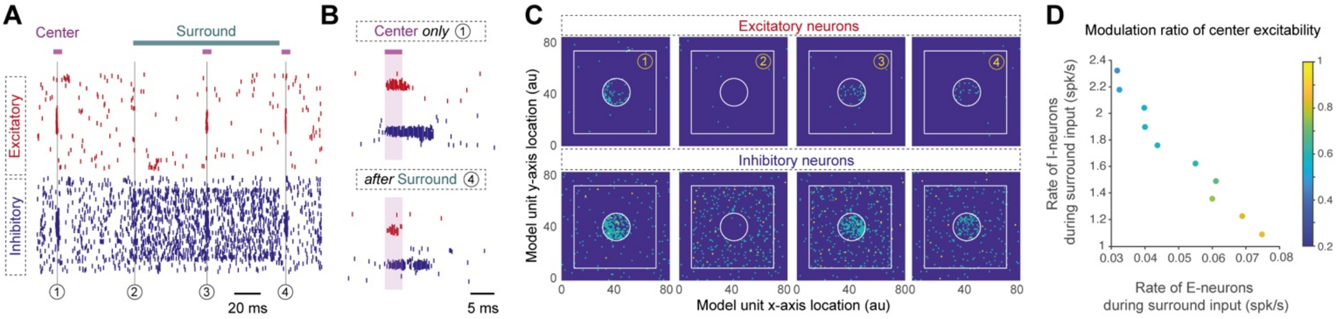

**Fig. S7. Complete raster of spiking network simulation (supplemental to Fig. 4)**

*(A)* Network model simulation using E/I connectivity patterns summarized in Fig. 4A. Top: Raster of spiking activity from 6400 excitatory and 6400 inhibitory neurons with four stimulus conditions that are marked with vertical numbered lines that simulate experimental conditions shown in Fig. 1. Conditions 1: center only before surround, 2: surround only, 3: center during surround, and 4: center after surround.

*(B)* Expanded network response of conditions 1 (top) and 4 (bottom) shown in (A).

*(C)* Spatial arrangement of center and surround network response profiles during the 4 different stimulus impulse conditions. The square schematically shows the region in which neurons were placed on a square grid. Excitatory (top) and inhibitory neurons (bottom) within the white circle received inputs corresponding to the center stimulation. All the neurons in white square but outside the inner circle received inputs corresponding to the surround network. Each subplot shows activity measured within a bin of 5 ms.

*(D)* Spiking frequency of model inhibitory neurons as a function of the spiking frequency of excitatory neurons. The inverse relationship is correlated of the peak amplitude ratio of the 'center during surround' divided by the 'center-only' response (*see* color map).

**Supplementary Table 1. Viruses used in this study**

| Name | Type | Titer | Source |
|---|---|---|---|
| rAAV2/hsyn-hChR2(H134R)-EYFP-WPRE-PA | AAV2 | $5.6 \times 10^{12}$ vg/mL | UNC Vector Core |
| rAAV2/hsyn-hChR2(H134R)-mCherry-WPRE-PA | AAV2 | $5.1 \times 10^{12}$ vg/mL | UNC Vector Core |
| rAAV2-EF1α-DIO-hChR2(H134R)-EYFP | AAV2 | $4.4 \times 10^{12}$ vg/mL | UNC Vector Core |
| ssAAV-2/2-shortCAG-dlox-hChR2(H134R)_EYFP(rev)-dlox-WPRE-hGHp(A) | AAV2 | $1.4 \times 10^{12}$ vg/mL | ZNZ Viral Vector Facility |
| rAAV2/hsyn-mCherry | AAV2 | $5.3 \times 10^{12}$ vg/mL | UNC Vector Core |
| rAAV5-CAG-FLEX-tdTomato | AAV5 | $7.8 \times 10^{12}$ vg/mL | UNC Vector Core |
| AAV5-EF1a-DIO-TVA-V5-t2A-RG | AAV5 | $1.2 \times 10^{12}$ vg/mL | DMC laboratory |
| EnvA-ΔG-Rb-EGFP | Modified Rabies | $1 \times 10^{1º}$ vg/mL | DMC laboratory |

**Supplementary Table 2. Drugs used in electrophysiology experiments**

| Compound | Drug name | Concentration | Source |
|---|---|---|---|
| Gabazine (SR 95531) | 6-Imino-3-(4-methoxyphenyl)-1(6*H*)-pyridazinebutanoic acid hydrobromide | 10 µM | Tocris Bioscience, USA |
| TTX | (4*R*,4a*R*,5*R*,7*S*,9*S*,10*S*,10a*R*,11*S*,12*S*)-Octahydro-12-(hydroxymethyl)-2-imino-5,9:7,10a-dimethano-10a*H*-[1,3]dioxocino[6,5-*d*]pyrimidine-4,7,10,11,12-pentol | 1 µM | Tocris Bioscience, USA |
| 4-AP | 4-Aminopyridine | 100 µM | Sigma-Aldrich |
| QX314 | *N*-(2,6-Dimethylphenylcarbamoylmethyl)triethylammonium bromide | 3 mM | Tocris Bioscience, USA |

**Supplementary Table 3. Stimulus input**

| Type of input | Stimulated neurons | Spike input | Stimulus PSP | Repeats |
|---|---|---|---|---|
| Center Input | All neurons (exc. and inh.) in a circle with radius 10 grid points.<br><br>314 exc. neurons and 314 inh. neurons. | Each neuron received 10 synchronous spikes corresponding to the presentation of a flash like center stimulus. Arrival time of each synchronous spike event was taken from a uniform distribution U[0,1] ms. | 1.03 mV at -54 mV to exc. neurons<br><br>0.86 mV at -54 mV to exc. neurons | 20 |
| Surround input | All neurons (exc. and inh.) within a square of size 60 grid point centered at the center of the network. Neurons part of the center stimulus were excluded.<br><br>3286 exc. neurons and 3286 inh. neurons. | Poisson type spiking input with a spike rate of 100 Hz.<br><br>Surround stimulus was presented for 200 ms. | 2.58 mV at -54 mV | 20 |

**Supplementary Table 4. Neuron parameters**

| Parameter | Value | Description |
|---|---|---|
| $C_m$ | 200 pF | Membrane capacitance |
| $g_L$ | 20 nS | Leak conductance |
| $V_{th}$ | -54 mV | Spike threshold |
| $E_L$ | -70 mV | Reversal potential for leak conductance |

| | | |
|---|---|---|
| $t_{ref}$ | 2 ms | Refractory time |

**Supplementary Table 5. Synapse parameters**

| Parameter | Value | Description | Notes |
|---|---|---|---|
| $J_{ex}$ | 0.63 mV | EPSP amplitude for connection from external excitatory inputs | EPSP amplitude was measured at a holding potential of -54 mV |
| $J_{ix}$ | -0.3 mV | IPSP amplitude for connection from external inhibitory inputs | IPSP amplitude was measured at a holding potential of -54 mV |
| $J_{ee}$ | {0.13, 0.26} mV | EPSP amplitude for exc. → exc. connection | EPSP amplitude was measured at a holding potential of -54 mV |
| $J_{ei}$ | {0.11, 0.21, 0.32, 0.42, 0.53, 0.63} mV | EPSP amplitude for exc. → inh. connection | EPSP amplitude was measured at a holding potential of -54 mV |
| $J_{ie}$ | {-0.14,-0.17, -0.19, -0.22, -0.28, -0.3} mV | IPSP amplitude for inh. → exc. connection | IPSP amplitude was measured at a holding potential of -54 mV |
| $J_{ii}$ | -0.055 mV | IPSP amplitude for inh. → inh. connection | IPSP amplitude was measured at a holding potential of -54 mV |
| $tau_e$ | 1.0 ms | Time constant of excitatory synapses | This was the same for both exc. → exc., exc. → inh. synapses and external excitatory input synapses |
| $tau_i$ | 3.0 ms | Time constant of inhibitory synapses | This was the same for both inh. → exc., inh. → inh. synapses and external inhibitory input synapses |
| $d_{ee}$ | 0.2 ms | Synaptic delay for exc. → exc. synapses | |

| | | |
|---|---|---|
| $d_{ei}$ | 0.1 ms | Synaptic delay for exc. → inh. synapses |
| $d_{ie}$ | 0.1 ms | Synaptic delay for inh. → exc. synapses |
| $d_{ii}$ | 0.1 ms | Synaptic delay for inh. → inh. synapses |

**Supplementary Table 6. Network connectivity**

| Parameter | Value | Description | Notes |
|---|---|---|---|
| $N_e$ | 6400 | Excitatory neurons | |
| $N_i$ | 6400 | Inhibitory neurons | |
| Excitatory grid size | 80 × 80 | Neurons were placed on a regular grid | |
| Inhibitory grid size | 80 × 80 | Neurons were placed on a regular grid | |
| $C_{ee}$ | 0.135 | Connection probability between exc. to exc. neurons. This number refers to the integral of the connectivity kernel | Based on our experimental data (*See* Fig. 3L) |
| $C_{ei}$ | 0.112 | Connection probability between exc. to inh neurons. This number refers to the integral of the connectivity kernel | Based on our experimental data (*See* Fig. 3L) |
| $C_{ie}$ | 0.315 | Connection probability between inh to exc. neurons. This number refers to the integral of the connectivity kernel | Based on our experimental data (*See* Fig. 3L) |
| $C_{ii}$ | 0.3375 | Connection probability between inh. to inh. neurons. This number refers to the integral of the connectivity kernel | Based on our experimental data (*See* Fig. 3L) |
| $S_e$ | 16 grid points | Standard deviation of Gaussian used to estimate distance dependent connectivity from exc. to exc. and exc. to inh. neurons. | |

| | | |
|---|---|---|
| | 2% of the network size | |
| $S_i$ | 20 grid points<br>2.5% of the network size | Standard deviation of Gaussian used to estimate distance dependent connectivity from inh. to exc. and inh. to inh. neurons. |

**Supplementary Table 7. External Input**

| Parameter | Value | Description |
|---|---|---|
| nu_exc | 1000 Hz | Poisson type spike trains were injected in all neurons |
| nu_inh | 600 Hz | Poisson type spike trains were injected in all neurons |
